# Mgl2^+^ cDC2 triggering of fungal allergic inflammation depends on a spore induced glycolytic shift fuelled by local availability of glucose

**DOI:** 10.1101/2025.07.13.664342

**Authors:** Julio Furlong-Silva, George Vere, Phuong Tuyen Nguyen, Nigel Gotts, Yun Xu, Stefano A.P. Colombo, Daniel P. Conn, Bethany L. McCann, Ana Cruz, Laila Alsharq, Craig Beall, Rafael Argüello, Shizuo Akira, Selinda Orr, Salomé LeibundGut-Landmann, Andrew S. MacDonald, Elaine Bignell, Gordon Brown, Darius Armstrong-James, Royston Goodacre, Peter C. Cook

**Affiliations:** Medical Research Council Centre for Medical Mycology at the University of Exeter, Department of Biosciences, UK; Centre for Metabolomics Research, Department of Biochemistry, Cell and Systems Biology, Institute of Systems, Molecular and Integrative Biology, University of Liverpool, UK; Lydia Becker Institute of Immunology and Inflammation, University of Manchester, Manchester, UK; Department of Clinical and Biomedical Sciences, University of Exeter Medical School, Exeter, UK; Faculty of Medicine, Department of Infectious Disease, Imperial College London, UK; Centre d’Immunologie de Marseille-Luminy, France; Laboratory of Host Defense, WP1 Immunology Frontier Research Center, Osaka University, Osaka, Japan; Wellcome-Wolfson Institute for Experimental Medicine, School of Medicine, Dentistry and Biomedical Science, Queen’s University Belfast, Belfast, UK; Section of Immunology, Vetsuisse Faculty and Institute of Experimental Immunology, University of Zürich, 8057 Zurich, Switzerland

## Abstract

Fungal spores are a major cause of severe asthmatic disease. However, the precise events that cause individuals to become sensitised to spores are poorly understood. Mgl2^+^ type 2 conventional dendritic cells (Mgl2^+^ cDC2s) are critical in coordinating allergic airway inflammation in mice following repeated exposure to inhaled spores. Yet, whether these DCs are directly acquiring spores from the airway, and the downstream mechanism(s) upon fungal uptake causing DCs to trigger allergic inflammation are unknown. Here we find that spores are acquired by lung DCs after inhalation although these events are rare (∼ 0.5% of the cDC2 population). Transcriptomics on isolated spore^+^ Mgl2^+^ cDC2s, compared to spore^-^ Mgl2^+^ cDC2s from the same environment, revealed that a major consequence of fungal uptake was a boost in metabolic activity. Single-cell metabolic profiling revealed this increase in Mgl2^+^ cDC2 metabolism upon spore acquisition was fuelled by a glycolytic shift. To pinpoint if nutrient availability and acquisition is an important determinant of this response, mass spectrometry-based metabolomics revealed that, during fungal allergic inflammation, the local airway nutrient environment is altered. To ascertain which of these could be fuelling DC responses, we identified which substrates that feed into glycolysis were crucial. Of these, we found that glucose availability acts as a key rheostat in shaping cDC2 responses to spores. These data highlight a crucial role for glycolytic metabolism in driving cDC2 responses to spores, which is governed by glucose availability, defining novel targets for future therapeutic development of fungal allergic inflammation.

## Introduction

Fungi are major, yet often underappreciated, triggers of allergic inflammation that underpins asthma^1^. Spores of the environmental mould *Aspergillus fumigatus* (*Af*), designated as a critical pathogen of concern by the World Health Organisation, are one of the primary fungi that cause these responses^2,3^. Of the 300 million individuals that suffer from asthma worldwide, approximately 10% patients display severe disease^4^. It is projected that 30-50% of these severe asthmatic patients are sensitive to fungi, which correlates with reduced treatment efficacy and poorer patient outcomes^5^. Despite this, these patients are difficult to diagnose and treat effectively^6^. Our limited understanding of the immune mechanisms responsible for triggering these harmful responses to spores hampers our ability to develop new approaches to diagnose and treat fungal allergic disease.

Allergic inflammation in asthma is underpinned by type 2 (e.g. IL-4, IL-5 and IL-13) and/or type 17 (e.g. IL-17 and IL-22) cytokine responses which orchestrate influxes of activated granulocytes, airway hyper-responsiveness (AHR), mucus production and fibrosis^7–9^. Studies have shown that both CD4^+^ T cells and innate lymphoid cells (ILCs) can be critical sources of type 2 and type 17 cytokines in response to fungal allergens^7,10^. We and others have found repeatedly exposing mice to *Af* spores causes fungal allergic airway inflammation that is dependent on type 2 and type 17 cytokine from CD4^+^ T cells^11–15^. We further found that dendritic cells (DCs), particularly a conventional subset of DCs (cDC2s) expressing Mgl2 (macrophage Gal/GalNAc lectin-2), were vital for triggering CD4^+^ T cell Th2 fungal allergic inflammation^11^. Furthermore, in humans we have identified (via sampling sputum) that the cDC2s in the airway is a key defining feature of patients with severe asthma and fungal sensitivity^16^. However, despite defining the primary DC subset responsible for fungal allergic inflammation, the mechanism(s) governing the ability of these cDC2s to trigger allergic type 2 responses in response to fungi has not yet been addressed.

Though DCs are integral for triggering allergic inflammation, it is unclear whether they directly encounter inhaled spores. Epithelial cells, airway macrophages (MΦ), neutrophils, DCs and monocytes have all been shown to acquire spores and co-ordinate protective immunity^17–22^. However, these findings were mostly in the context of invasive fungal disease (aspergillosis) whereby mice were either immunosuppressed prior to *Af* infection and/or exposed to a large fungal inoculum (typically 10 - 100 million spores) resulting in a dramatically different inflammatory environment compared to conditions in healthy individuals or in allergic disease^19,23–25^. Due to their small size (2-3 μM) spores are likely deposited deep in alveolar spaces, even in healthy immunocompetent individuals^26^. Mathematical modelling proposes airway MΦs clear inhaled spores^27,28^, but the role of DCs and whether frequent exposure to fungi impacts this process is unknown. Previous studies suggest that DCs likely encounter fungi, we have shown that early growth stages of spores promote DCs to trigger allergic inflammation, and others have shown that DC expression of pathogen recognition receptors C-type lectin receptors (CLRs) and Toll-like receptors (TLRs) is an important step in coordinating antifungal immunity^29–33^. However, it is also possible that other signals from neighbouring cell types encountering fungi may be more important in conditioning DCs to induce allergic inflammation. For example, cytokines released from epithelium cells (e.g. IL-33 and TSLP) upon contact with allergens and/or ILC secretion of cytokines (e.g. IL-13 and leukaemia inhibitory factor), promote DC induction of allergy^34–37^. Therefore, to decipher how cDC2s co-ordinate fungal allergic inflammation, the question arises if cDC2s directly encounter fungi in the airway and whether this an important step in cDC2 activation.

In this work, we administered fluorescent spores to mice in a model of direct, intranasal (i.n.) sensitisation and challenge with repeated low doses of live *Af* conidia to induce allergic inflammation and tracked the uptake of spores. Few spores were acquired by the epithelium, with the majority acquired by immune cells. Furthermore, upon repeat spore exposure the uptake of fungi by cDC2s increased after repeated administration, although these remained rare events (only ∼100 lung cDC2s acquired spores). Combining the use of transcriptomic analysis on isolated DCs with cytometry-based SCENITH metabolic profiling, we show that fungal uptake boosts metabolic activity, fuelled by glycolysis, when compared to cDC2s from the same environment which had not acquired spores. We also show that CLR-Card9 signalling helps promote the increase of cDC2 metabolic activity and activation upon spore uptake. Mass spectrometry-based metabolomics revealed that fungal allergic inflammation alters the nutrient environment of the airway and compound profiling determined that cDC2s rely on nutrients that feed into glycolysis. Importantly, of the identified nutrients, only glucose was essential, and its availability is crucial in determining the magnitude of cDC2 responses upon spore exposure. Overall, use of complementary approaches reveals that cDC2s, which are vital for driving downstream fungal allergic inflammation, uptake of spores induces activation that is reliant on increased glycolysis and is governed by glucose availability.

## Results

### Immune cells acquire spores during fungal allergic inflammation

The precise host cell types (e.g. airway epithelial versus immune cells) which coordinate removal of inhaled spores in healthy individuals is unclear. To address this directly, we exposed mice i.n. to a single dose of surface labelled *Af* spores (4 x 10^5^) and following 3h, administered CD45-APC antibody i.t. immediately prior to lung harvest to identify airway immune cells^38^. Staining precision cut lung slices (PCLS) with EpCAM and podoplanin, to visualise the epithelial network^39,40^, revealed most of the spores were acquired by airway CD45^+^ immune cells, with only a small proportion of *Af* associated with the epithelium and/or outside cells in the airways (Fig. 1A-C and Supplementary Fig. 1). Having identified that immune cells were responsible for spore uptake, flow cytometry analysis after a single exposure of GFP-expressing *Af* (1x *Af*) revealed spores were largely acquired by Alv MΦs, with only minimal uptake by other immune cell populations (Fig. 1E & F and Supplementary Figs. 2-4). Therefore, Alv MΦs are primarily responsible for acquiring *Af* following a single exposure to fungal spores.

**Figure 1.**
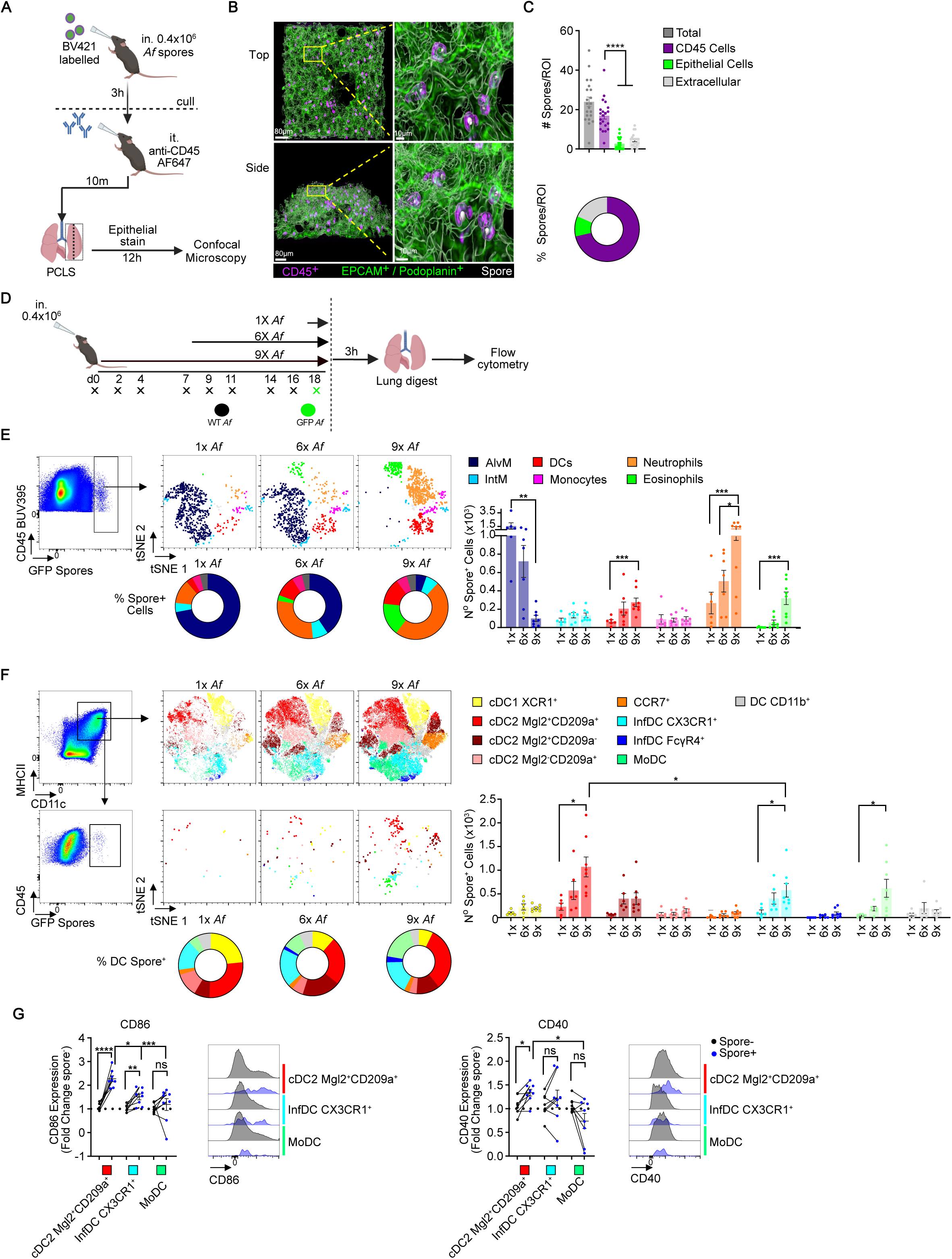
Repeated exposure to fungi increases cDC2s acquisition of fungal spores. **A**) Mice were exposed to labelled *A. fumigatus* (*Af*) spores (strain: CEA10, 4 x 10^5^ spores) via intranasal (i.n.) transfer, 3h later following CD45 administration i.n. lung tissue was harvested for Precision Cut Lung Slices (PCLS). **B**) Representative images show PCLS from top and side views illustrating spores (white), airway CD45^+^ spore^+^ immune cells (purple) or epithelial cells (green) of *Af*-exposed mice. **C**) Graphs display the number and proportion of spores in airway CD45^+^ cells, epithelium or in extracellular spaces from PCLS of *Af*-exposed mice. ROI: region of interest. **D**) Naïve mice or mice that were repeatedly exposed to unlabelled *Af* via intranasal transfer on the indicated time points, were exposed to GFP-expressing *Af*. Tissues were harvested 3h after the first, sixth or ninth dose of spores (1x, 6x and 9x, respectively). **E**) Representative tSNE plots reveal the immune cell populations that acquire spores by unbiased clustering analysis of flow cytometry data from the lung of mice exposed to a single or repeat dose of spores. Graphs display the numbers and percentages of spores acquired by alveolar macrophages (AlvM), interstitial macrophages (IntM), DCs, monocytes, eosinophils and neutrophils from the lung tissue. **F**) Representative tSNE plots reveal the DC subsets that acquire spores by unbiased clustering analysis of flow cytometry data from the lung of mice exposed to *Af* spores. Graphs display the numbers and percentages of spores acquired by DC subsets from the lung tissue. **G**) Representative flow cytometry overlay plots identify expression of CD40 and CD86 on spore^+^ and spore^-^ DCs from the lungs of mice exposed to 9 doses of spores. Graphs show the fold change in expression of spore^+^ versus spore^-^ DCs of each DC subset. The 3 DC subsets which acquire the greatest number of spores are shown. **A** – **C**) Number of spores quantified from Region of Interest (ROI) pooled from 3 independent experiments (6-8 ROIs per experiment, 1 mouse per experiment), *n* = 3 biologically independent animals). **D** – **G**) data are from two independent experiments (n = 6 - 8 biologically independent animals per group). Significance was determined by **B**) an unpaired 2-tailed Student’s t-test between indicated groups, **E** & **F**) 1-way ANOVA with Tukey’s multiple-comparison post-hoc test or **G**) a paired Student’s t test *P < 0.05, **P<0.01, ***P < 0.001, **** P<0.0001, NS not significant,

We and others have found that exposing mice repeatedly to spores causes allergic inflammatory responses^11,12^. To identify whether Alv MΦ are still the dominant population to acquire spores following the onset of fungal allergic inflammation, we exposed mice repeatedly to 6 or 9 doses of spores, which induces hallmarks of allergic inflammation (Supplementary Fig. 2). Replacing the final dose with GFP^+^ spores revealed constant *Af* exposure resulted in a dramatic reduction in the number of spore^+^ Alv MΦs (Fig. 1D & E and Supplementary Figs 2-4). This decline in spore uptake by Alv MΦs was not compensated by either lung interstitial MΦs or monocytes previously implicated in mediating microbe killing and inflammation^41,42^, as both populations expanded but the number of spore^+^ Int MΦs and monocytes did not increase (Fig. 1E and Supplementary Figs. 2-4). Instead, we found following 6x and 9x doses of spores, neutrophils, eosinophils and DCs were the major lung cell populations to increase their spore uptake (Fig. 1E and Supplementary Figs. 2-4). These alterations in spore uptake did not change the total number of spores acquired in the lung as numbers of spore GFP^+^ cells remained similar regardless of whether mice received a single or multiple *Af* doses (Supplementary Fig. 4A).

These data show that the dynamics of spore uptake and clearance is fundamentally altered in the context of constant spore exposure, characterised by increased uptake by granulocytes and DCs.

### Mgl2^+^ cDC2s are the major subset to acquire spores during fungal allergic inflammation

After identifying that lung DC acquisition of spores increases after repeatedly dosing mice with *Af*, we wanted to ascertain if certain DC subsets are preferentially acquiring *Af*, as we have previously demonstrated that one of these (Mgl2^+^ cDC2s) are crucial for mediating fungal allergic inflammation^11^. In mice that received only a single dose of GFP^+^ *Af*, so in the absence of an allergic inflammatory environment, ∼100 total DCs acquired spores of which cDC2s were the largest majority (Fig. 1F and Supplementary Figs. 5 & 6). In agreement with our previous findings, constant exposure to spores caused the expansion of: cDC2s (Mgl2^+^CD209a^+^, Mgl2^+^CD209a^-^, Mgl2^-^CD209a^+^), CCR7^+^ DCs, monocyte derived DCs (MoDCs), CX3CR1^+^ inflammatory DCs (InfDCs) while others (cDC1s, FcγR4 cDCs) remained static or decreased (Supplementary Figs. 5 & 6A). Of these, only Mgl2^+^CD209a^+^ cDC2s, Inf DCs and moDCs increased their uptake of spores as allergic inflammation developed (9x versus 1x) and cDC2s represented the dominant subset to acquire spores, accounting for 45-50% of total DC spore^+^ cells (Fig. 1F, Supplementary Figs. 5 & 6A). However, even upon allergic inflammation DC spore uptake in the lung remained relatively rare as only ∼1% of cDC2s acquired spores (∼100 spore^+^ cDC2s per lung; Supplementary Fig. 6A).

To determine whether spore acquisition induced DC activation, we measured expression of co-stimulatory molecules CD40 and CD86, pivotal in polarising CD4^+^ T cells to secrete type 2 cytokines^43,44^, on DCs from *Af*-exposed mice. Of the DC subsets to acquire fungi after 9x doses of *Af*, only spore^+^ Mgl2^+^CD209a^+^ cDC2s significantly upregulated (versus spore^-^ Mgl2^+^CD209a^+^ cDC2s from the same lung tissue) both CD40 and CD86 expression (Fig. 1G and Supplementary Fig. 6B). In contrast, the increase in CD86 expression on spore^+^ CXCR3^+^Inf DCs was more modest (Fig. 1G and Supplementary Fig. 6B).

Taken together, these data show that Mgl2^+^CD09a^+^ cDC2s are the dominant lung DC subset to acquire spores, accompanied with increased activation of these DCs compared to neighbouring cDC2s and other DC subsets in the same environment.

### Spore acquisition, not the lung environment, is a more critical driver of altered Mgl2^+^CD209a^+^ cDC2 gene expression

Having established that we could effectively identify lung DCs that had acquired spores from the airway, we wanted to determine the functional impact of *Af* acquisition by Mgl2^+^CD209a^+^ cDC2s. Though we, and others^11,37,45–48^, have revealed that cDC2s are effective in triggering allergic inflammation, whether and how direct spore acquisition or the lung environment governs cDC2 ability to trigger allergic inflammation is unknown.

To define the impact of spore acquisition versus the lung environment on cDC2s during fungal allergic airway inflammation, we employed RNA-sequencing on spore^+^ and spore^-^ Mgl2^+^CD209a^+^ cDC2s isolated from the lungs of mice exposed to 6x or 9x *Af* doses alongside Mgl2^+^CD209a^+^ cDC2s isolated from control (PBS exposed) mice (Fig. 2A). Principal Component Analysis (PCA) of the 500 most expressed genes revealed PC1 (∼50%) and PC2 (∼23%) separated samples based on *Af* spore uptake and number of exposures, respectively. This resulted in spore^+^ cDC2s from both 6x and 9x *Af* dosed mice clustering separately from spore^-^ cDC2s which were isolated from the same tissue environment, with minor, but markedly less separation occurring between spore^-^ and PBS control dosed samples (Fig. 2B). Together, these data suggest that while gene expression of cDC2s is influenced by the inflammatory signals in the lung environment, direct spore uptake plays a much more dominant role on the transcriptome of cDC2s during allergic inflammation.

**Figure 2:**
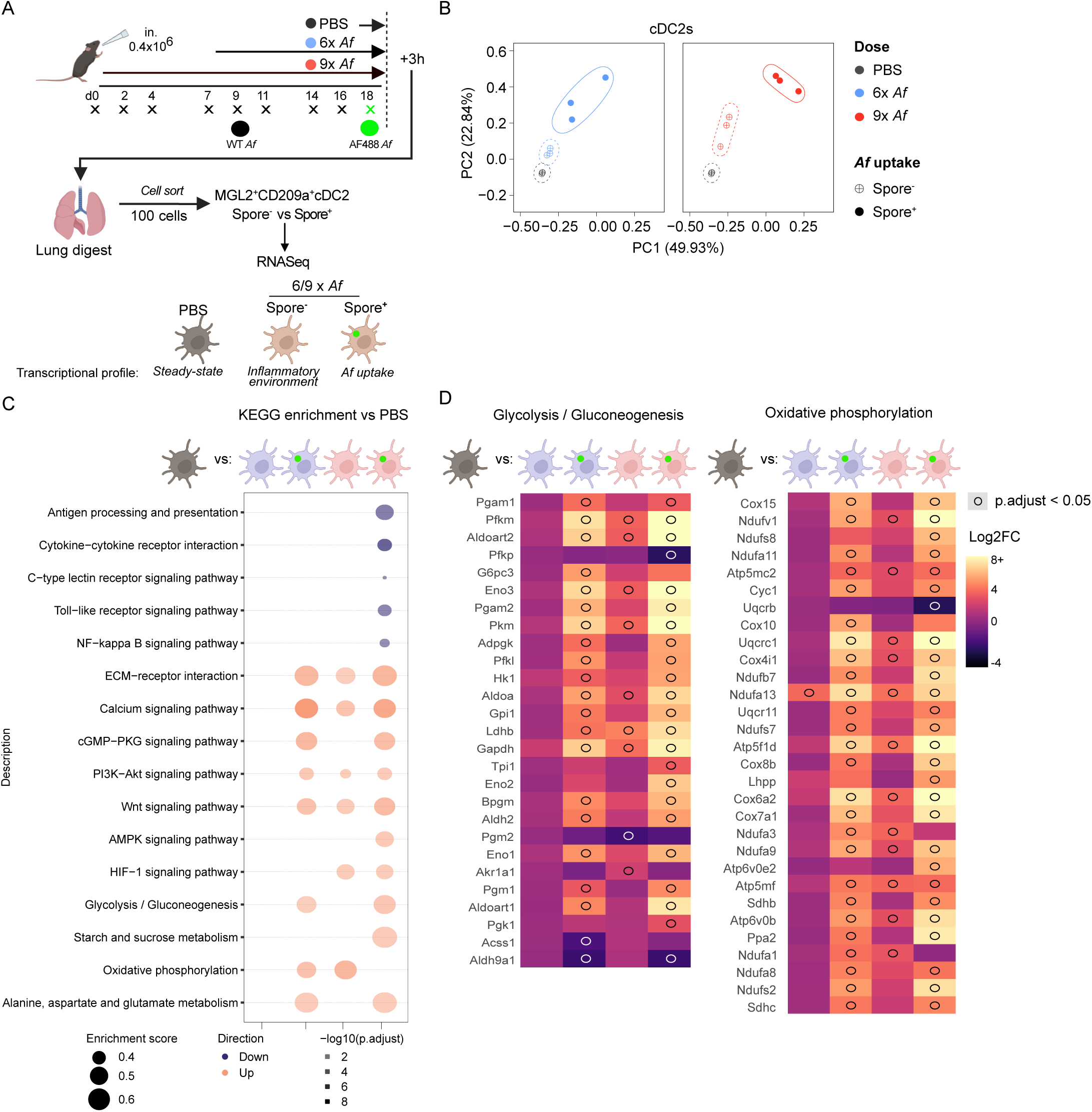
Spore acquisition, not the lung environment, causes a greater change of gene expression within cDC2s. **A**) Mice were repeatedly exposed to unlabelled *Af* spores (4 x 10^5^ per dose) via intranasal for 5 or 8 doses followed by a final dose of labelled *Af*. An additional control group of mice received 9 doses of PBS only. Tissues were harvested 3h post sixth and ninth dose of spores (6x and 9x, respectively). Spore^-^ and spore^+^ Mgl2^+^CD209a+ cDC2s were isolated from the lung tissue of mice that received 6x and 9x doses of *Af*, alongside Mgl2^+^CD209a^+^ cDC2s from the lung tissue of PBS exposed mice. Transcriptome of each population was generated using low input kits followed by RNA sequencing (Illumina). **B**) Principal Component Analysis (PCA) of mRNA transcript expression of spore^-^ (crossed) and spore^+^ (solid) Mgl2^+^CD209a^+^ cDC2s isolated from 6x (blue) and 9x (ref) *Af*-exposed mice or Mgl2^+^CD209a^+^ cDC2s isolated from PBS treated mice (grey). Plotted circles represent 95% confidence interval of indicated groups. Each dot represents an individual pooled sample. **C)** KEGG pathway analysis show significant up or down-regulated transcriptional pathways of indicated spore^+/-^ Mgl2^+^CD209a^+^ cDC2s from *Af*-exposed mice versus Mgl2^+^CD209a^+^ cDC2s from PBS mice. **D**) Heatmaps show scaled gene expression of genes associated with glycolysis/gluconeogenesis and oxidative phosphorylation of indicated spore^+/-^ Mgl2^+^CD209a^+^ cDC2s from *Af*-exposed mice versus Mgl2^+^CD209a^+^ cDC2s from PBS mice. Circles indicate p.adjust < 0.05 compared to PBS controls calculated via Wald test in DESeq2. Each individual sample is a pool from 3-4 biologically independent animals derived for 3 independent experiments.

To determine the major gene pathways within cDC2s that are shaped by spore uptake versus the allergic lung environment, we undertook KEGG pathway enrichment analysis. As revealed via PCA analysis, a greater number of identified pathways were up and down-regulated from spore^+^ cDC2s compared to spore^-^ cDC2s in *Af*-exposed mice (Fig. 2C, Supplementary Fig. 7 & 8 and Supplementary Data 1). Unexpectedly, cDC2 gene pathways associated with fungal recognition (especially CLRs^49,50^) were either unaltered or down-regulated upon spore acquisition (Fig. 2C and Supplementary Figs.7 & 8). Gene pathways that have been associated with DC induction of allergic inflammation that includes antigen presentation/TFs/NFκB pathways (*Irf4*^46^, *Mbd2*^51^), cytokine receptors (*Il4ra*^52,53^, *Il13r1*^37^), chemokines (*ccl17*, *ccl22*^51^) and co-stimulatory molecules (*cd40*^43^, *cd86*^44^, *Icosl*^54^ and *Pdcd1lg2*^55^) were either decreased or unaltered upon spore uptake by cDC2s (Fig. 2C and Supplementary Fig. 8). Instead, spore^+^ cDC2s had elevated expression of genes associated with cellular migration that is associated with the induction of allergic inflammation^56^ (Supplementary Fig. 8). However, the most prominent changes when comparing cDC2s from *Af*-exposed mice (both spore^+^ and spore^-^), to cDC2s from PBS treated mice, were upregulation of signalling pathways associated with metabolic activity (e.g. calcium, PI3K-Akt and HIF-1 signalling) (Fig. 2C and Supplementary Figs. 7 & 8). Comparing spore^+^ and spore^-^ cDC2s from *Af*-exposed mice revealed several metabolic gene pathways involved with glycolysis (including fructose and mannose metabolism), oxidative phosphorylation (OXPHOS), fatty acid degradation and glycerolipid metabolism were increased (Fig. 2C & D and Supplementary Figs. 7 & 8).

In summary, these data revealed for the first time that only cDC2s which acquire spores, during fungal allergic inflammation undergo widespread metabolic reprogramming.

### The response of cDC2s to spores is dependent on glycolysis

To verify that spore acquisition by cDC2s during fungal allergic inflammation enhances metabolic activity, we utilised SCENITH (Single-Cell ENergetIc metabolism by profiling Translation inHibition)^57–60^ on spore^+/-^ cDC2s from the lung tissue of *Af* exposed mice (Fig. 3A-C). Spore^+^ cDC2s had greater puromycin incorporation than spore^-^ cDC2s from the same lung environment, indicating that *Af* uptake has increased their translation rate and therefore overall metabolic activity (Fig. 3B & C). Treatment with metabolic inhibitors (2-deoxyglucose, DG, and/or Oligomycin A, O) revealed that cDC2s were completely reliant on glucose/glycolysis and not Fatty Acid Oxidation (FAO) (Fig. 3D). Furthermore, spore acquisition enhanced cDC2 glycolytic capacity, concurrent with a reduction in mitochondrial dependency, when compared to spore^-^ cDC2s (Fig. 3D). These data confirm our gene expression analysis revealing that the increase of Mgl2^+^CD209a^+^ cDC2 metabolic activity upon spore uptake is fuelled by glycolysis.

**Figure 3.**
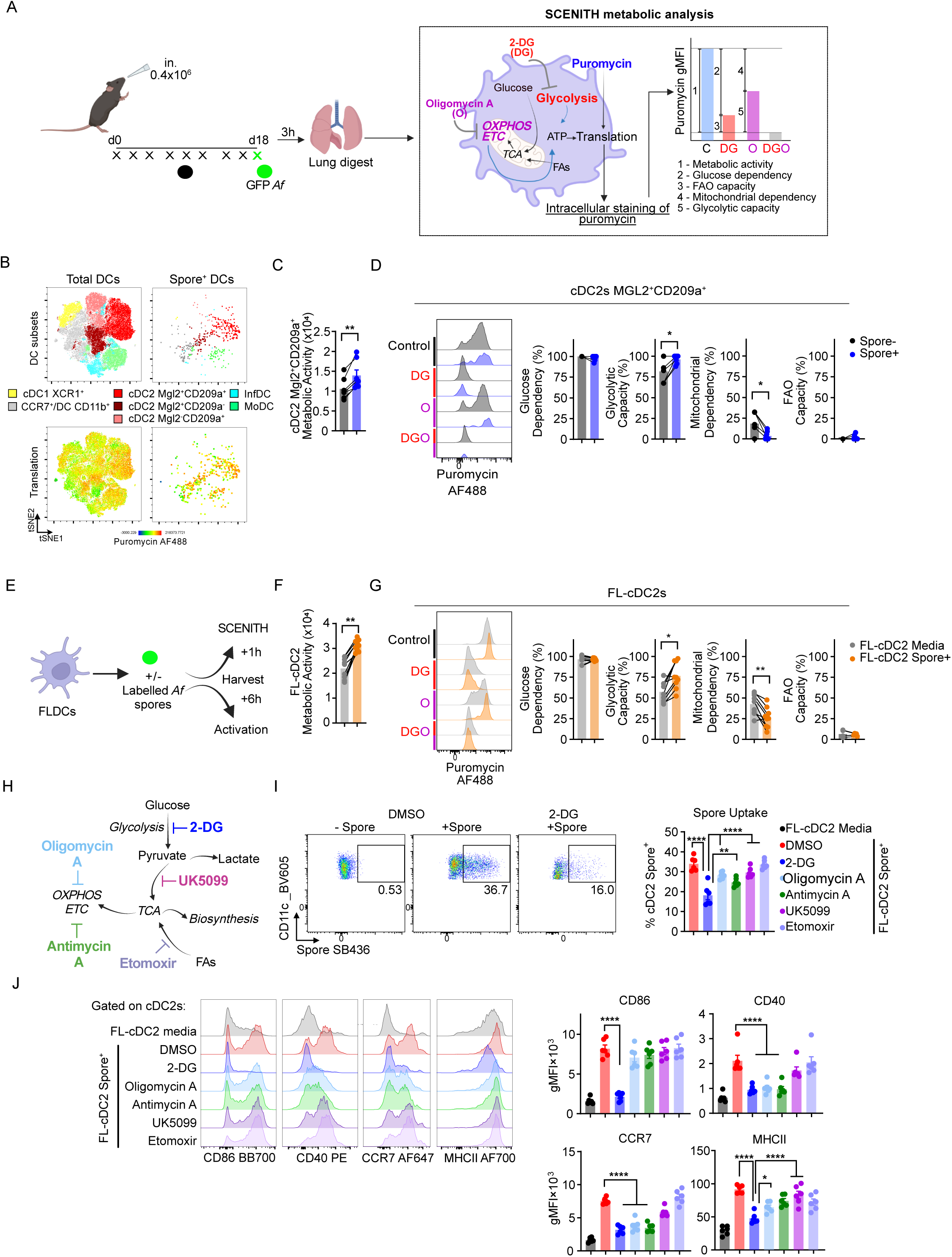
Spore acquisition by cDC2s and downstream activation is dependent on glycolysis. **A**) Mice were repeatedly exposed to unlabelled *Af* spores (4 x 10^5^ per dose) i.n. transfer for 8 doses then received a final dose of labelled *Af*. Lung tissue was harvested 3h after the ninth dose of spores (9x) and processed for SCENITH metabolic analysis via flow cytometry (left). Principle of SCENITH single cell metabolic profiling; puromycin staining is a measure of cellular translation. Each sample is independently treated ex vivo with control (Co), 2-DG, Oligomycin-A or 2-DG + Oligomycin, gMFI of puromycin staining of each treatment indicates metabolic activity, glucose dependency, glycolytic capacity, mitochondrial dependency and fatty acid oxidation (FAO) capacity). (right) **B**) Representative tSNE plots reveal the DC subsets that acquire spores and staining of puromycin (indicates rate of translation) by unbiased clustering analysis of flow cytometry data from the lungs *Af*-exposed mice. **C**) Graph displays metabolic activity (as detailed in **B**) of spore^+/-^ Mgl2^+^CD209a^+^ cDC2s from the lungs of Af-exposed mice. Connecting lines indicates populations from the same single animal. **D**) Representative flow cytometry overlay plots of gMFI puromycin staining of spore^+/-^ Mgl2^+^CD209a^+^ cDC2s from Af-exposed mice that were treated *ex vivo* as detailed in **B**). Graphs display the corresponding metabolic dependency and capacity parameters of spore^+/-^ Mgl2^+^CD209a^+^ cDC2s. **E**) Flt3L bone marrow derived DCs (FLDCs) generated from mice were exposed to labelled 4-hour swollen Δ*pyrG Af* spores and processed for SCENITH after 1h, or for measuring DC activation after 6h incubation. **F**) Graph displays metabolic activity (as detailed in **B**) of spore^+^ FL-cDC2s after 1h of exposure to labelled *Af* or untreated FL-cDC2s (media). Connecting lines indicates FL-cDC2 populations derived from a single animal. **G**) Representative flow cytometry overlay plots of gMFI puromycin staining of spore^+/-^ FL-cDC2s exposed to labelled *Af* that were treated as detailed in **D**). Graphs display the corresponding metabolic dependency and capacity parameters of spore^+^ FL-cDC2s after 1h of exposure to labelled *Af* or untreated FL-cDC2s (media). **H**) Several inhibitors were utilised to disrupt different metabolic pathways. **I**) Representative flow cytometry plots of FL- cDC2 uptake of labelled spores in the presence and absence of 2-DG for 6h. Graphs display FL-cDC2 uptake of spores following exposure to labelled *Af* in the presence of separate inhibitors for 6h. **J**) Representative flow cytometry overlay plots and graphs display show the gMFI of co-stimulatory molecules (CD40 and CD86), CCR7 and MHCII on FL-cDC2s after exposure to spores in the presence or absence of separate inhibitors. **A** - **D**) data are from two independent experiments (n = 7 biologically independent animals per group). **E** – **K**) data from either three independent (E & G) or two independent experiments (I & J) (*n*= 6 or 9 FLDCs derived from biologically independent animals per group. Significance was determined by 2-tailed Student’s t-test (**C**,**D**,**F** & **G**) or one-way ANOVA with Tukey’s multiple-comparison post-hoc test (**I** & **J**) between indicated groups. *P < 0.05, **P<0.01, ***P < 0.001, **** P<0.0001.

Though spore uptake appears to trigger enhanced metabolic activity of cDC2s (Figs. 2 and 3A-D), the allergic inflammatory environment (such as type 2 cytokines) in *Af*-exposed mice may be involved in conditioning cDC2s to trigger glycolysis upon spore exposure. For example, upon exposure to type 2 cytokines MΦ increase both glycolysis and OXPHOS^61–64^. Therefore, to determine whether spore uptake by itself is sufficient to boost glycolytic activity, we utilised *in vitro* Flt-3 ligand bone marrow derived DC (FLDCs) which generate cDC1, cDC2 and plasmacytoid DCs (pDC) like subsets^65^ (Supplementary Fig. 9). Stimulating FLDCs with Δ*pyrG* spores (an auxotrophic *<u>Af</u>* strain requiring uracil/uridine supplementation to grow, enabling generation of swollen but arrested growth stages^29^) caused greater FL-cDC than FL-pDC activation with increased expression of costimulatory molecules. (CD40, CD86), CCR7 (crucial for DC migration) and MHC-II accompanied with release of proinflammatory cytokines and chemokines (Supplementary Fig. 9). To test whether spore stimulation impacts FLDC metabolism, we could not use arrested Δ*pyrG* spores as we discovered that the Δ*pyrG* spores are themselves metabolically active via extracellular flux analysis, with evident extracellular acidification rate (ECAR) and oxygen consumption rate (OCR) (Supplementary Fig. 10A). Instead, we treated FLDCs with killed swollen spores (which induced similar FLDC activation (Supplementary Fig. 10B) causing a rapid increase in both ECAR and OCR, demonstrating that exposure to spores alone is sufficient to boost DC metabolic activity (Supplementary Fig. 10C). To determine if a certain FLDC subset was predominantly responsible for this shift in metabolic activity, we first identified that that FL-cDC2s had the greatest capacity to acquire fungi versus other FLDC subsets (Supplementary Fig. 11). Furthermore, SCENITH analysis revealed spore^+^ FL-DC2s had boosted metabolic activity compared to untreated (media) FL-DC2s which was entirely reliant on glucose dependency and glycolytic capacity (Fig. 3E-G). Taken together, these data mirror what we had observed with *in vivo* cDC2s from the lung and confirm spore exposure, independent of allergic inflammation, is a key trigger of DC2 activation and enhanced glycolysis.

To determine the crucial metabolic pathways fuelling DC activation in response to fungi, we exposed FLDCs to labelled spores in the presence of metabolic inhibitors targeting glycolysis (2-DG), OXPHOS (Oligomycin A and Antimycin A), the TCA cycle (UK5099) and FAO(Etomoxir) (Fig. 3E & H). Inhibition of glycolysis via 2-DG dramatically reduced spore uptake, FL-cDC2 activation and secretion of IL-12p40 and CCL22 (Fig. 3I, J and Supplementary Fig. 12A). Treatment of 2-DG had no impact on *Af* germination, demonstrating that only DC pathways were affected (Supplementary Fig. 12B). Inhibition of all other metabolic pathways only mildly impacted FL-cDC2 spore uptake, activation and antigen presentation, with the magnitude of any reductions lower than those observed under glycolysis inhibition (Fig. 3I & J).

These data demonstrate that glycolysis plays a critical role in fuelling cDC2 responses to fungi including spore uptake, downstream activation, migratory potential and antigen presentation.

### The response of cDC2s to spores are partially dependent on CLR signalling

After identifying that spore uptake is a crucial step in mediating glycolysis dependent activation of cDC2s, we wanted to understand if CLR or TLR signalling pathways underpinned these responses. Though our RNAseq dataset did not show upregulation of these pathways upon spore uptake (Fig. 2 and Supplementary Fig. 8), other studies have highlighted these pathways can be critical in co-ordinating immunity against *Af* and other non-fungal allergens^30–32^. To investigate the role of the CLR signalling of *Af*-induced activation of DCs, we first tested the role of Card9, a downstream signalling adaptor of most CLRs^33^ implicated in mediating fungal allergic inflammation^66^. The ability of FL-cDC2s generated from *Card9*^-/-^ mice to acquire swollen spores was similar to WT FL-cDC2s (Fig. 4A & B). Furthermore, expression of CD40, CD86, CCR7 and MHC-II of spore^+^ WT and *Card9*^-/-^ FL-cDC2s was significantly increased compared to untreated (media) *Card9*^-/-^ FL- cDC2s (Fig. 4C). However, spore^+^ *Card9*^-/-^ FL-cDC2 expression of CD40 and CCR7 was significantly lower than spore^+^ WT FL-cDC2s suggesting a partial role of CLR signalling in controlling cDC2 responses. Similar results were obtained when FLDCs were treated with a Syk inhibitor (R406), to blockade CLR signalling^33^, during spore exposure, which decreased FL-cDC2 expression of activation markers (Supplementary Fig. 13B). To determine the impact of individual CLRs on FL-cDC2 activation, we tested the impact of Dectin-1, Dectin-2 and/or Mincle deficiency on coordinating DC responses to spores (Supplementary Fig13A). Individual deficiency of either Dectin-1 or Dectin-2 had no impact on the ability of cDC2s to upregulate surface expression of activation markers (Supplementary Fig 13B). To test whether individual CLRs may display functional redundancy for *Af* induced cDC2 activation, we generated FL-DCs deficient in Dectin-1 and Dectin-2 double knockout (DKO) or Dectin-1, Dectin-2 and Mincle triple knockout (TKO). In response to spores, DKO and TKO FL-cDC2 expression of activation markers (CD40, CD86 and MHCII) was significantly reduced compared to WT FL-cDC2s (Supplementary Fig. 13B), while no difference was observed between DKO and TKO Fl-cDC2s. In contrast, to test the role of TLR signalling, we found that FL-cDC2s generated from *Myd88*^-/-^ and *Trif*^-/-^ mice mounted similar responses to that of FL-cDC2s from WT mice (Supplementary Fig. 13B). These data highlight the importance of the CLR pathway in co-ordinating cDC2 activation in response to *Af* spores.

**Figure 4.**
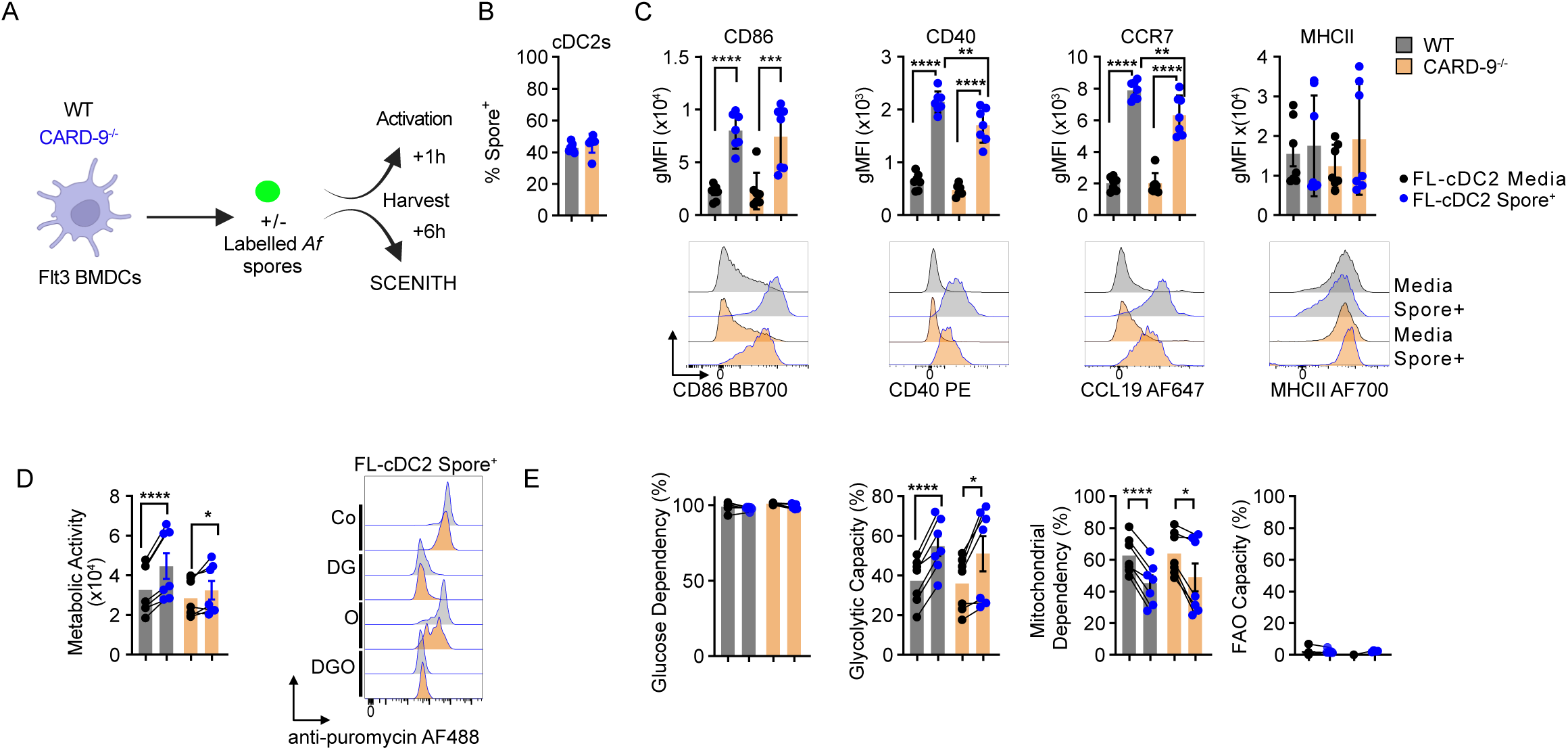
Card9-dependent signalling in response to spores is involved in activating cDC2s. **A**) Flt3L bone marrow derived DCs (FLDCs) generated from WT or *Card9*^-/-^ mice were exposed to labelled 4-hour swollen Δ*pyrG Af* spores and processed for SCENITH after 1h, or for measuring DC activation after 6h incubation. **B**) Graphs display WT and *Card9*^-/-^ FL- cDC2 uptake of spores following exposure to labelled *Af*. **C**) Representative flow cytometry overlay plots and graphs display show the gMFI of co-stimulatory molecules (CD40 and CD86), CCR7 and MHCII on WT and *Card9*^-/-^ FL-cDC2s after exposure to spores or untreated (media). **D & E**) Representative flow cytometry overlays and graphs show metabolic activity, dependency and capacity of WT and *Card9*^-/-^ FL-cDC2s after 1h of exposure to labelled spores or untreated (media). Connecting lines indicates FL-cDC2 populations derived from a single animal. Data are from two independent experiments (*n* = 7 biologically independent animals per group). Significance was determined by one-way ANOVA with Tukey’s multiple-comparison post-hoc test (**C)**, or a paired Student’s t-test (**E**) between indicated groups. *P < 0.05, **P<0.01, ***P < 0.001, **** P<0.0001.

To test whether CLR pathways are implicated in triggering the increase of glycolysis by FL- cDC2s in response to *Af* we utilised SCENITH on *Card9*^-/-^ FLDCs exposed to labelled spores. The overall metabolic activity of both WT and *Card9*^-/-^ spore^+^ FL-cDC2s increased compared to media only treated FL-cDC2s, however the increase observed in spore^+^ *Card9*^-/-^ FL-cDC2s was more muted compared to spore^+^ WT FL-cDC2 counterparts (Fig. 4D). This increase in both WT and *Card9*^-/-^ FL-cDC2s activity was fuelled by glycolysis (Fig. 4E), showing that CLR disruption did not alter the metabolic pathways which fuel DC activation.

These data demonstrate that CLR signalling plays an important role in controlling the ability of cDC2 to respond to spores and are partially responsible for fuelling the critical glycolytic shift necessary to support these responses.

### Alteration of the local airway nutrient environment during fungal allergic inflammation

The extracellular tissue environment is known to be a critical regulator of innate immune cell metabolism controlling their responses in a variety of disease settings. We have previously shown that the lung airway nutrient availability regulates glycolysis and OXPHOS metabolic activity of alveolar macrophages, shaping their ability to respond to type 2 inflammation^38^. Therefore, we hypothesized that the local availability of nutrients in the airway during fungal allergic inflammation may be a further crucial signal that governs the critical glycolytic boost that occurs within cDC2s upon uptake of spores.

To determine if fungal allergic inflammation alters the availability of nutrients within the airway, we employed LC-MS-based metabolomics on broncho-alveolar lavage (BAL) fluid and serum samples collected from spore exposed or control mice (Fig. 5A). PCA of the top 500 metabolite features in BAL samples revealed PC1 (∼32% both positive and negative mode) and PC2 (∼11.2% positive mode, 13.2% negative mode) clearly separated samples from *Af*-exposed versus control mice (Fig. 5B & C). In contrast, PCA on metabolites in the serum showed no separation in samples from *Af*-exposed versus control mice. This reveals that only the lung nutrient environment is altered upon fungal allergic inflammation with no changes systemically.

**Figure 5.**
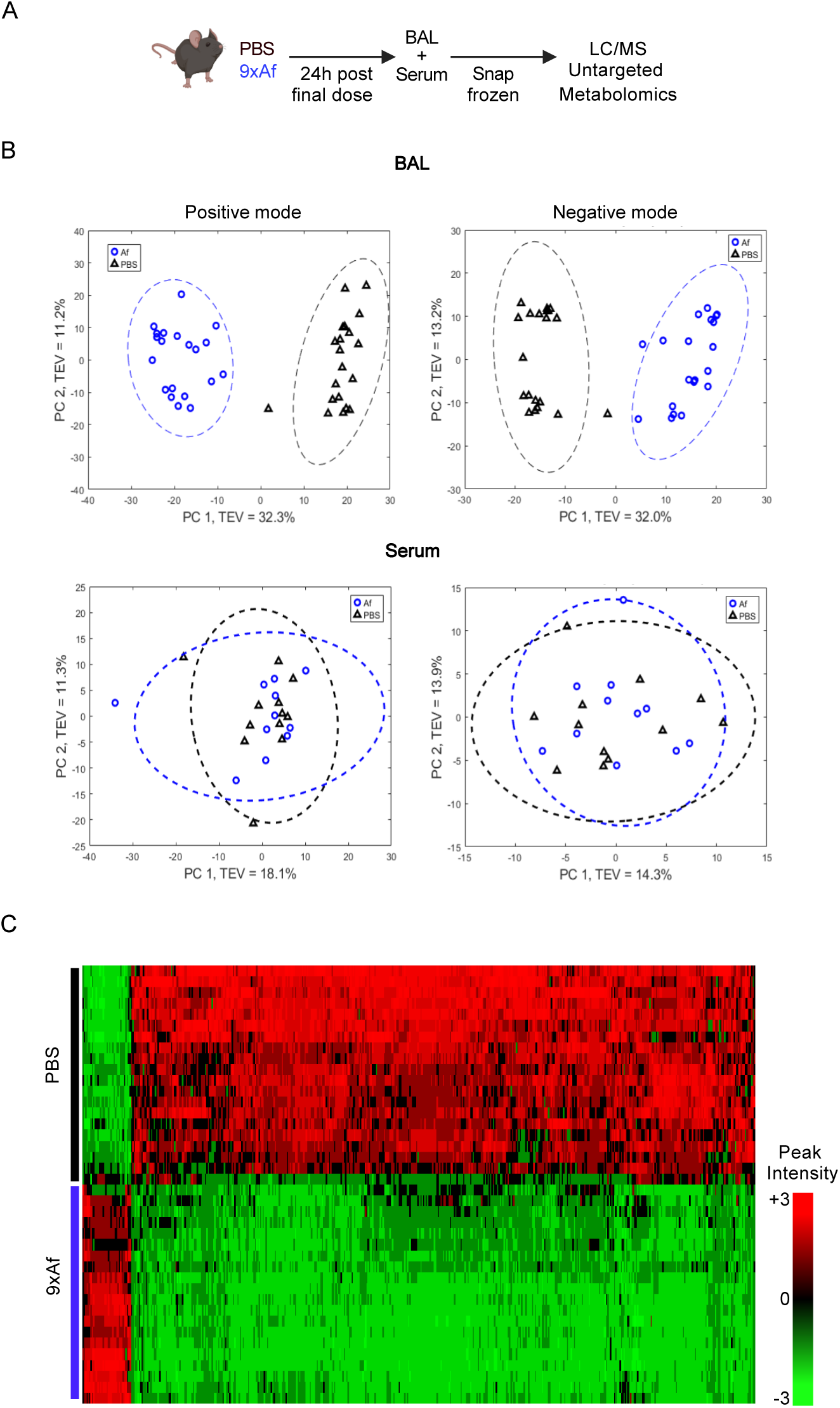
Fungal allergic inflammation alters the local airway nutrient environment. **A**) Mice were repeatedly exposed to unlabelled *Af* spores (4 x 10^5^ per dose) or PBS i.n. for 9 doses. Broncho-Alveolar lavage (BAL) fluid and serum were harvested 24h after the last dose of spores. Samples from *Af*-exposed mice (9x*Af*; blue circles) or PBS (black triangles) were processed for untargeted Liquid Chromatography- Mass spectrometry (LC-MS) metabolomics **B**) PCA score plots derived from untargeted LC-MS metabolomics from BAL (top) and serum (bottom) isolated from *Af* or PBS exposed mice. Plotted circles represent 95% confidence intervals of relevant groups. **C**) Heatmap of positive mode top 500 LC-MS features with FDR <0.05. Data are from four independent experiments (*n* = 20 biologically independent animals per group).

As the local airway environment is altered, it could be an important regulator of observed glycolysis that occurs within cDC2s after spore uptake.

### Local availability of glucose governs cDC2 spore responses

The shift in the airway nutrient environment in repeat *Af*-dosed mice is also accompanied with large influxes of inflammatory cells which mediate allergic responses (Supplementary Fig. 2)^11^. Utilisation of SCENITH on cells isolated from the lungs of *Af*-exposed mice revealed that the influx in many of these inflammatory cell types, especially eosinophils which are the largest population to be recruited to the airway (Supplementary Fig. 14), are highly metabolically active and display a highly glycolytic profile. This suggests that spore^+^ cDC2s (much rarer than these inflammatory influxes) may be competing for key glycolytic related metabolites with much larger populations of equally glycolytic immune populations within the same nutrient environment. To pinpoint the crucial nutrients in the airway, which are impacted during fungal allergic inflammation, it is essential to determine which nutrient sources are specifically utilised by cDC2s to fuel activity upon spore acquisition. To address this, we exposed FLDCs to spores and employed Biolog’s Phenotype MicroArray Mammalian plates PM-M1 that are pre-loaded with an array of individual carbon nutrients known to be utilised in both glycolysis and OXPHOS metabolism (Fig. 6A). This approach has been previously employed to identify inosine utilisation by T effector cells^67^ and glycogen by DCs^68^. We found that FLDC activity upon spore exposure is fuelled by nutrients supporting glycolysis, notably glucose and glycolysis related intermediary nutrients that includes glucose-6-phosphate, fructose-6-phosphate, glycerol-3-phosphate and mannose (Fig. 6B). Adenosine and inosine, two purine nucleotides, were also capable of supporting FLDC metabolism which may be derived from the cleavage of the ribose sugar to support central carbon metabolism^67^ Notably, neither galactose (a primary nutrient utilised in oxidative phosphorylation^69–71^), nor pyruvate or TCA intermediates (downstream of glycolysis^72^) could sustain FLDC metabolism (Fig. 6B).

**Figure 6.**
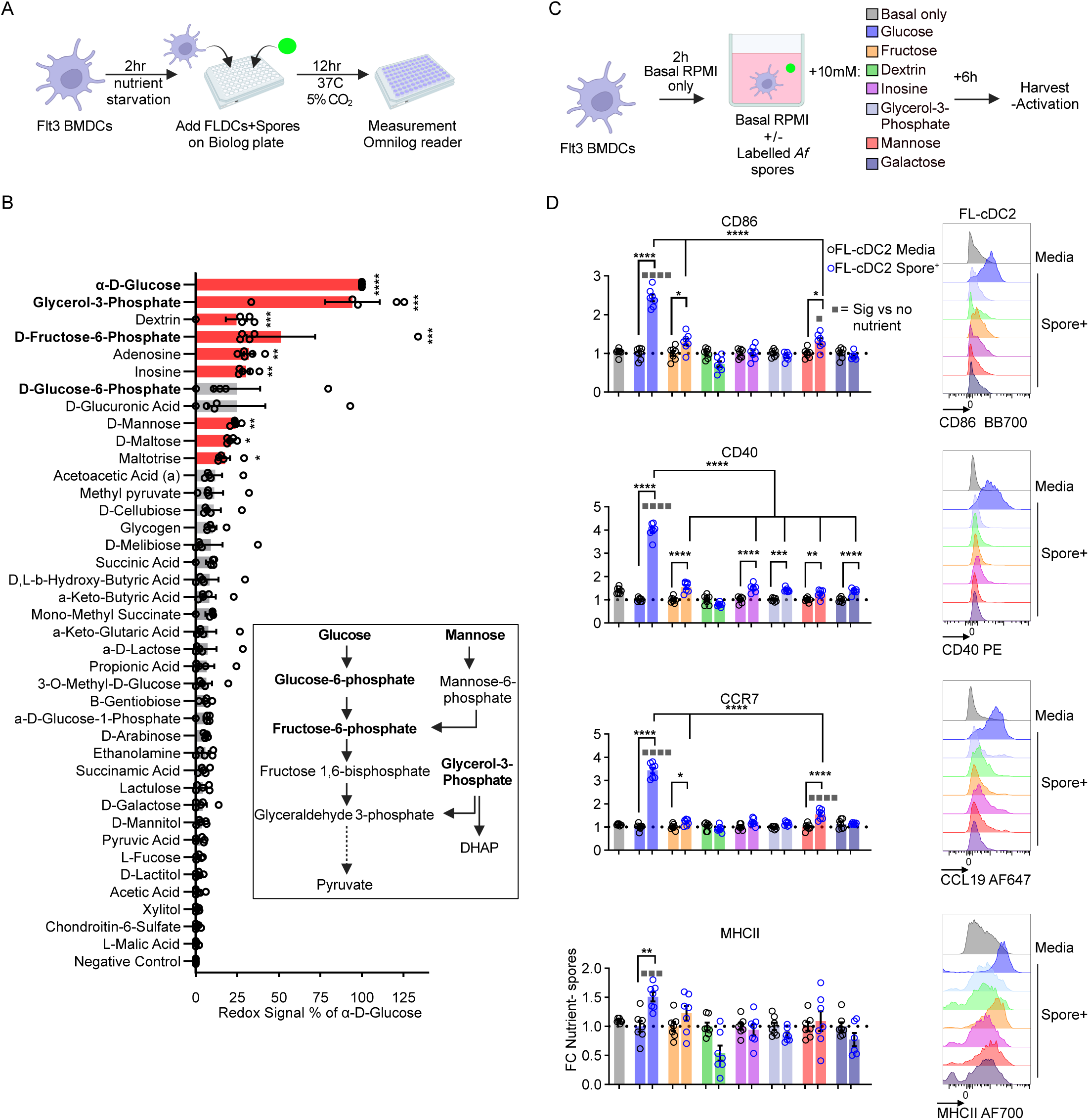
Carbon sources, especially glucose, are essential for cDC2 responses to fungal spores. **A**) Flt3L bone marrow derived DCs (FLDCs) generated from mice were exposed to 4-hour PFA-killed spores on Biolog M1 Carbon Source Plates with metabolic activity measured with addition of redox dye. **B**) Graph displays FLDC redox signal in culture with PFA-killed spores when incubated on each compound, which is normalised to the percentage of activity detected when cultured in α-D-Glucose. Compounds which caused significant metabolic activity of FLDCs compared to negative control are coloured in red. Schematic shows the points in which identified nutrients can feed into glycolysis. **C**) FLDCs generated from mice were precultured in basal media prior to addition of spores in the presence of specified nutrients for 6h. **D**) Representative flow cytometry overlays display the gMFI of co- stimulatory molecules (CD40 and CD86), CCR7 and MHCII on FL-cDC2s c after exposure to spores whilst individually supplemented with glucose, fructose, dextrin, inosine, glycerol-3-phosphate, mannose or galactose. Graphs display the fold change difference in expression compared to FL-cDC2s not exposed to spores in basal media. **B)** data from two independent experiments (*n* = 5 FLDCs derived from biologically independent animals). **D**) data from two independent experiments (*n* = 7 FLDCs derived from biologically independent animals). Statistical significance determined by **B**) Kruskal Wallis ANOVA with Dunn’s multiple-comparison post-hoc test vs negative control or **D**) one-way ANOVA with tukey’s multiple comparison test between indicated groups. *P < 0.05, **P<0.01, ***P < 0.001, **** P<0.0001.

To determine which of these identified nutrients are essential in fuelling cDC2 spore responses, we cultured FLDCs with spores in basal media, selectively supplemented with identified nutrients (Fig. 6B & C). Of the nutrients tested, only the addition of glucose fully restored cDC2 ability to both acquire spores and upregulate expression of activation markers (CD40, CD86, CCR7 and MHC-II) (Fig. 6C, D and Supplementary Fig. 15). Supplementation of other tested nutrients did not support FL-cDC2 activation, demonstrating that cDC2s rely on glucose as the major nutrient source to fuel their glycolytic shift in response to fungal spores.

As fungal allergic inflammation alters metabolites in the lung airway, and cDC2s rely on glucose for activation to spores we wanted to ascertain if glucose concentrations from the BAL fluid from PBS or *Af*-exposed mice were impacted (Fig. 7A). We found that BAL glucose concentrations were decreased in the BAL fluid from *Af*-exposed mice compared to PBS- exposed mice (Fig. 7B). This decrease is likely reflective of influxes of a large number of glycolytic active cells (e.g. eosinophils) into the lung thereby increasing local glucose consumption (Supplementary Figs. 2 & 14). However, the glucose concentrations we detected in the BAL fluid from both *Af*-exposed and PBS-exposed mice was substantially lower than levels present in the blood (∼5 mM)^73,74^. Therefore, to determine whether these lower concentrations of glucose present in the airway is a factor in sustaining cDC2 responses to spores, we cultured FLDCs in basal media supplemented with varying amounts of glucose starting from 10mM (typical concentrations used in standard tissue culture media, e.g. RPMI) (Fig. 7C). As expected, cultures of FLDCs in glucose concentrations of 10mM (RPMI) and 5mM (blood) fuelled spore^+^ FL-cDC2 activation (expression of CD40, CD86, CCR7 and MHC-II) following spore exposure (Fig. 7D). However, when incubated in 0.4mM glucose (reflecting healthy lung airways in humans^75,76^) spore^+^ FL-cDC2s were significantly less activated (Fig. 7D). Reducing glucose concentrations below 0.4mM further impacted FL- cDC2s activation in response to spores. These observed changes were not due to an alteration in FL-cDC2 viability or their ability to acquire spores (Supplementary Fig. 16).

**Figure 7.**
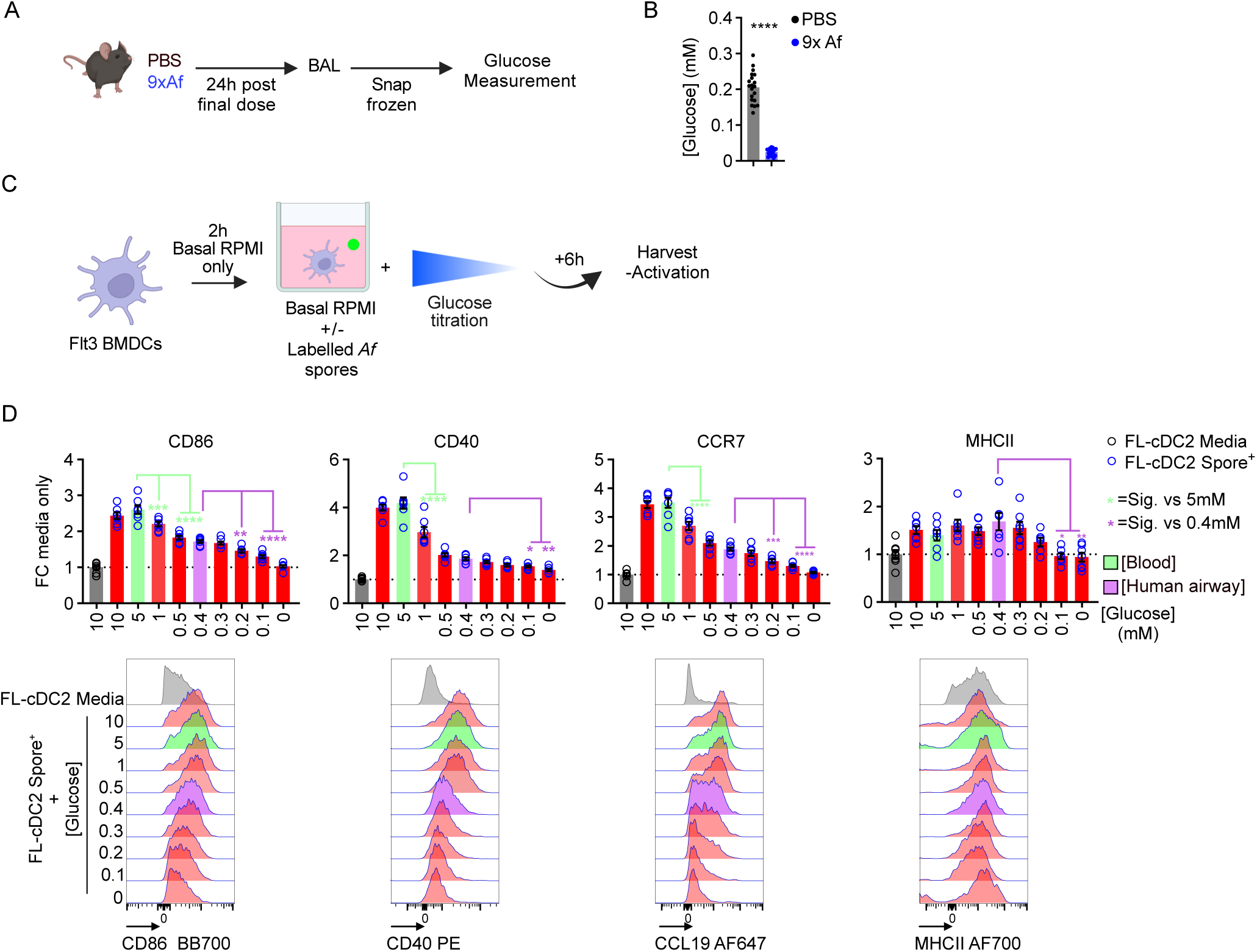
Availability of glucose governs the capacity of cDC2s to respond to fungal spores. **A**) Mice were repeatedly exposed to unlabelled *Af* spores (4 x 10^5^ per dose) or PBS i.n. for 9 doses. Broncho-Alveolar lavage (BAL) fluid was harvested 24h after the last dose. **B**) Graph displays glucose concentrations in the BAL fluid from *Af* or PBS exposed mice. **C**) Flt3L bone marrow derived DCs (FLDCs) generated from mice were precultured in basal media prior to addition of labelled 4-hour swollen Δ*pyrG Af* spores in the presence of different concentrations of glucose for 6h. **D**) Representative flow cytometry overlays display the gMFI of co-stimulatory molecules (CD40 and CD86), CCR7 and MHCII on FL-cDC2s after exposure to spores whilst individually supplemented with different concentrations of glucose. Graphs display the fold change difference in expression compared to FL-cDC2s not exposed to spores in basal media. **B)** Data are from four independent experiments (*n* = 20 biologically independent animals per group). **D**) Data are from two independent experiments (*n* = 8 FLDCs derived from biologically independent animals). Statistical significance determined by **B**) two-tailed student’s t-test or **D**) one-way ANOVA with tukey’s multiple comparison test between indicated groups. *P < 0.05, **P<0.01, ***P < 0.001, **** P<0.0001.

Together, these data show that despite widespread nutrient changes upon allergic inflammation, availability of glucose acts is the crucial rheostat in fuelling cDC2 glycolysis, shaping their responses to fungal spores.

## Discussion

Using strategies to track the acquisition of fungal spores in the airway, along with complementary single cell technologies, we have identified that cDC2s, which trigger fungal allergic inflammation, undergo metabolic reprogramming characterised by increased glycolysis upon spore uptake. We further reveal that levels of nutrients in the airway are altered during fungal allergic inflammation, with glucose availability fuelling the glycolysis enabling cDC2s to respond to spores.

Previous studies propose the lung airway epithelial cells acquire inhaled fungi and/or allergens releasing inflammatory mediators (e.g. IL-33, IL-25, TSLP and GMCSF) to promote immunity^17,24,36,77–80^. However, these findings were predominantly drawn from either fungal infection models (with immunocompromised mice and/or a large inoculum) or allergy models with allergen extracts^20,77,78,81,82^. By developing an approach to track *Af* in the lung after administering a low number of spores into immunocompetent mice, we found that immune cells, not the epithelium, were critical for acquiring fungi (Fig. 1). Confirming previous modelling^27,28^, we found that Alv MΦ were the dominant immune cell population to initially acquire spores, though a small number of spore^+^ granulocytes and DCs were also identified. This changed dramatically after the onset of allergic inflammation with an increased number of spore^+^ DCs and granulocytes. Though the number of monocytes and Int MΦs increased upon constant spore exposure, here we found that few acquired spores. Though both populations have been associated with co-ordinating antifungal immunity and allergy^18,19,41,42^, the fact that few acquired *Af* implied these populations are not directly involved in causing fungal allergic inflammation. These approaches highlight that tracking the cell types which directly acquire allergens/pathogens reveals greater insight in determining the key orchestrators of allergic inflammation.

We have recently discovered that a particular cDC2 subset, that express Mgl2, are critical for initiating type 2 fungal allergic inflammation^11^. Tracking fungal uptake across all lung DC subsets showed the majority were acquired by Mgl2^+^ cDC2s, and that this increased DC activation compared to neighbouring spore^-^ Mgl2^+^ cDC2s (indicated via CD40 and CD86 expression) (Fig 1). While inflammatory and monocyte derived DCs have been implicated in antifungal immunity^19^, fewer of them acquired spores, and the ones that had appeared less activated compared to spore^+^ Mgl2^+^ cDC2s. This highlights that tracking spore uptake is effective at indicating the DCs which are likely triggering downstream inflammation. To uncover mechanisms by which cDC2s mediate allergic inflammation, some approaches have been focused on isolating cDC2s from the lung and rely on downstream transcript analysis (via RNAseq or scRNAseq), typically comparing to cDC2s from control lung tissue (e.g. naïve)^11^. However, here we find spores^+^ cDC2s, which would go onto stimulate downstream inflammation, is rare with only ∼100 spore^+^ Mgl2^+^ DCs (∼1% of cDC2s). Therefore, it is possible previous studies detected genes that are important for maintaining the function of the whole cDC2 population, but not specifically the ones which acquire antigen. Indeed, this may explain why transcript analysis of spore^+^ vs spore^-^ cDC2s revealed that numerous pathways associated with DC induction of type 2 inflammation (*Irf4*^46^, *Mbd2*^51^, *Il4ra*^52,53^, *Il13r1*^37^, *il2*^83^, *ccl17/ccl22*^51^, *Pdcd1lg2*^55^) were not up-regulated upon spore acquisition (Fig. 2). Further work is needed to understand the role of previously established pathways within cDC2s and whether they regulate responses upon acquisition of spores or other allergens that mediate type 2 inflammation.

Undertaking transcript analysis on spore^+^ and spore^-^ cDC2s from the same lung revealed that spore uptake had a significant impact on gene expression (Fig. 2). This revealed that one of the major impacts of fungal acquisition by cDC2s, which via single cell metabolic profiling we found was underpinned with increased glycolysis (Fig. 3). Previous studies have highlighted the crucial role of glycolysis to enable functional activity of MΦs, monocytes and DCs in responding to microorganisms, airway allergic inflammation^84–88^. However, these previous studies have not considered the role of spore uptake versus other environmental signals in triggering these metabolic changes. To knowledge, we have shown for first time that fungal induced boosting of glycolysis, which is essential for cDC2 activation, only occurs in DCs that acquire spores and not neighbouring spore^-^ cDC2s. This extends our understanding of how cDC2s undertake energy intensive tasks such as spore antigen processing and migration.

CLR signalling is essential for mediating antifungal immunity^89–91^, and studies have shown CLR mediated modulation of glycolytic metabolism licences MΦ and monocyte antifungal responses^84,92^. Furthermore, studies showed β-glucan increased DC glycolysis via CLR-syk signalling^93,94^. Our data showed that CLR-syk signalling, while not critical for uptake of cDC2s, was partially involved in cDC2 increase glycolysis and activation upon spore acquisition (Fig. 4). The precise ligands on the spores which activate these responses is unclear. We have previously demonstrated spore sensing by DCs is enhanced following spore swelling and growth^29^, suggesting that exposure of CLR ligands upon spore swelling is important. Future work for the identification of these components will be critical to better understand how spore^+^ cDC2s mediate fungal allergic inflammation and what other innate signalling pathways maybe involved.

The lung environment has been shown to be a crucial regulator of innate immune cell activity^38^. One aspect in which this influences immune cell activity and microbial infections, is the altered profile of nutrients in the airway compared to those observed in circulation^73,95,96^. In other settings, such as cancer, studies have highlighted altered concentrations of nutrients inside tumours can reduce the effectiveness of recruited immune cells^97^. Yet, prior to this study, the role of nutrients in shaping DC activation to fungi and causing allergic inflammation was unclear. By performing metabolomics on lung BAL fluid of *Af*-exposed mice, screening DC nutrient usage, to inform supplementation of defined nutrients into the media revealed that cDC2s rely almost exclusively on glucose to fuel increased glycolysis in response to spores (Figs. 5 & 6). Furthermore, we found that the magnitude of DC activation was dependent on glucose availability, especially within the range of airway glucose concentrations which are present in the airway^73^. Maintaining low glucose levels in the airway has the benefit of restricting microbial growth^95,96^ and our findings suggest an additional advantage is that these also partially restrain the ability of DCs to mediate inflammatory responses. Conversely, when this is reversed in hyperglycaemic conditions DC metabolism is altered, leading to chromatic remodelling which increases susceptibility to viral infection^98^. Indeed, increased airway glucose levels has been reported in diabetic patients^76^, which has been associated with higher incidence of asthma^99–102^. Further work is needed to understand whether metabolic disorders and/or therapies which target glycolysis (e.g. Metformin) alter glucose levels in the airway and whether this could influence airway DC responses and susceptibility to respiratory disease.

In summary, our work highlights that by tracking the DC populations which acquired spores after inhalation we were able to interrogate the processes which govern cDC2 capability to mediate fungal allergic inflammation. In this way, our data provide further insight in to how metabolic activity and levels of nutrients co-ordinate DC driven lung inflammation that is relevant for understanding allergy that underpins asthma and other diseases.

## Materials and methods

### Experimental animals

C57BL/6 were either purchased from Charles River Laboratories or bred in-house along with: *Card9*^-/-^^103^ and littermate controls at the University of Exeter. Dectin-1^-/-^9292, Dectin-2^-/-^ ^90^, and Mincle^-/-^^104^ were bred at Charles River Laboratories, both Dectin-1^-/-^Dectin-2^-/-^ and Dectin-1^-/-^Dectin-2^-/-^Mincle^-/-^^105^ were kindly gifted by S.O. All knockout strains were bred on C57BL/6 background. All mice were maintained under Specific Pathogen Free conditions and experiments approved under a project license granted (to either G.D.B (P6A6F95B5) or P.C.C. (PP6094315)) by the Home Office UK in addition to the University of Exeter AWERB. Experiments were performed in accordance with the United Kingdom Animals (Scientific Procedures) Act of 1986. Aged and sex-matched male and female mice aged 6-20 weeks were used for FLDC generation and *in vivo* experiments.

### Mouse model of anti-fungal allergic inflammation

To emulate anti-fungal allergic inflammation, a murine mode of repeat *Aspergillus fumigatus* (*Af*) exposure was used as previously described^11,12^. Briefly, mice were intranasally administered 4x10^5^ spores (CEA-10) in 50 μl PBS 0.05%Tween 80 (PBST) or vehicle control on days 0, 2, 4, 7, 9, 11, 14, 16 and 18. Allergic inflammation was measured by euthanising mice 24 hours after dosing on either day 5, 12 or 19. When assessing spore uptake, mice were euthanised 3h post final dose on either day 0, 11 or 18.

### Generation and analysis of Precision-Cut Lung Slices (PCLS)

Mice administered spores as above, were euthanised 3 hours post spore dose and immediately administered 2μg APC conjugated anti-mouse CD45 antibody intratracheally. PCLSs were then generated as previously described with minor modifications117117. Briefly, lungs of relevant mice were inflated with 1 mL of 1.5% UltraPure Low Melting Point Agarose (ThermoFisher Scientific) in PBS and left to solidify. Inflated lungs were then dissected and 300 µm slices generated utilising a Vibrating-blade microtome VT1000S (Leica). PCLS’s were then fixed with 1% PFA in PBS at room temperature (RT) for 15-20 minutes, washed several times with PBS and incubated with specific primary antibodies overnight at 4°C in PBS. Stained PCLS slices were then mounted on glass slides, covered with a glass coverslip and stored at 4oC until undertaking confocal microscopy and image analysis. PCLS’s were imaged on a Spinning-disk confocal microscopy (Dragonfly), operating under Fusion software and using a Nikon 10x/20x objective. PCLS were first wholly mapped using 10x z-stacks set at 2 µm. Subsequently, 6-8 fields of views representative of the whole slice were defined and subsequent 20x z-stacks set at 1 µm intervals generated. Acquired z-stack images were then analysed by 3D rendering of spores, epithelial and CD45+ immune cells utilising IMARIS software (version10.2). Spores were rendered utilising IMARIS Spot function (Spot size 2-5 µm), whilst epithelial and immune cell rendering were generated utilising Surface function. Following rendering, numbers of spores and cells were detected and enumerated utilising IMARIS software. Number of spore positive epithelial and CD45+ immune cells were defined utilising IMARIS filter function with shortest distances between spores and cells set at values of 0 and below. Numbers of extracellular spores were defined and enumerated based on shortest distances between spores and cells, set at values above 0.

### Isolation of immune cells from BAL and lung

Airway immune cells were obtained from BAL in 3x0.5ml washes with PBS + 2% FCS + 2 mM EDTA (all sigma-referred to as lung buffer). Lungs were processed by placing lung tissue in 0.8 U/mL Liberase TL, 80U/mL DNase in HBSS (all Sigma) and processed using the GentleMACS protocol 37C_m_LIDK_1. Digestions were halted by adding ice-cold lung buffer, followed by maceration through a 70μM cell strainer and subsequent RBC lysis using RBC lysis buffer (Biolegend) as per manufacturer’s protocol. In experiments where SCENITH was undertaken, the SCENITH protocol steps preceded the RBC lysis step. Following a final wash in lung buffer, cells were utilised in downstream flow cytometry or cell sorting.

### *Aspergillus* spore culture and labelling

Spores were cultured as previously described^11^, briefly CEA-10^107^ or A1160 ΔpyrG^108^ were grown on Sabouraud Dextrose Agar (SAB, Oxoid), at 37 °C (supplemented with 5mM uridine and 5mM uracil (u/u, both Sigma) for growth of Δ*pyrG*). Swollen time-arrested spores were generated as previously described^29^, briefly, Δ*pyrG* spores were harvested and resuspended in RPMI (Sigma) at 1.6 x10^7^ spores/ml in the presence or absence of 5 mM uricil/uridine, followed by incubation at 37C for defined periods. Generated swollen spores were washed extensively with PBS containing 0.05% Tween-80 (PBST) (both sigma) and stored in RPMI1640 (sigma) at 4 °C for up to 24h prior to use. CEA-10 Spores utilised in *in vivo* infections were harvested and resuspended in PBST. Aliquots of concentrated spore suspension were stored at -80°C and for each dose, a fresh aliquot was diluted so 4 x 105 (in 50 μl) spores in PBST ready for intranasal administration to mice. In experiments were killed spores were utilised, spores were resuspended in 1% PFA(Sigma) overnight followed by extensive washing in PBST before final resuspension in RPMI1640.

Fluorescently labelled spores were generated as previously described^81^. Briefly, freshly harvested CEA-10 or Δ*pyrG* spores were incubated in 50mM carbonate buffer (pH 8.3) + 0.5 mg/mL Sulfo-NHS-LC-LC-Biotin (Thermofisher) for 1 h at RT under continuous rotation. Unbound Sulfo-NHS-LC-LC-Biotin was then removed by incubating spores with 100 mM Tris-HCl (pH 8) for 15 minutes and subsequently washed a further 2 times in PBS. Spores were then incubated with PBS + 1:100 Streptavidin (SA) - conjugated to either Brilliant Violet (BV) 421 (Biolegend) for *in vivo* experiments or SuperBright (SB) 436 (Thermofisher) for FLDC experiments for 1 h at RT. Spores were then washed with PBST prior to *in vivo* intranasal (i.n.) dosing or spore germination in media (FLDC experiments).

To generate killed Af spores for assays, spores were treated with 1% paraformaldehyde (PFA) dissolved in PBS overnight and subsequently washed 5 times with PBS.

### FLDC culture and assays

Flt3-L derived bone marrow dendritic cells (FLDCs) were generated as previously described^65^, briefly, harvested bone-marrow cells from relevant mouse strains were cultured in RPMI1640 Glutamax (sigma) + 15% Foetal Calf serum (Sigma) + 50 μM 2- mercaptoethanol (Invitrogen) + 200 ng/mL recombinant Flt3-L (PeproTech) for 8 days at 37C in 5% CO_2_. Following 8 days, generated FLDCs were plated in complete RPMI1640 Glutamax + 10% FCS (Sigma) + 50 ng/mL Flt3-L + 50 μM 2-mercaptoethanol and, dependent on experiment, stimulated with either spores at a multiplicity of infection (MOI) of 5:1 or 10 μg/mL Zymosan (Invivogen) for defined periods.

### FLDC metabolic manipulation and nutrient utilisation assays

In FLDC metabolic inhibitor experiments, generated FLDCs were pretreated with: 10mM 2-DG (0.5% PBS), 50μM Etomoxir (0.5% PBS), 1μM Oligomycin A (0.1% DMSO), 1μM - Antimycin A (0.1% DMSO) and 5μM UK5099 (0.1 DMSO) in RPMI 1640 Glutamax + 10% FBS- +50 ng/mL Flt3-L 50uM 2-mercaptoethanol (Complete media) (all reagents sigma) for 30 mins, followed by the addition of 4 hour swollen Δ*pryG* spores at a MOI of 5:1 for 6 hours at 37°C 5% CO_2_. Following incubation, FLDCs were washed thoroughly in PBS+ 2% FCS + 2μM EDTA (FACS buffer, all sigma) for onward analysis.

For glucose titration and nutrient supplementation experiments, FLDCs were washed twice in RPMI-1640 + L-Glutamine, No Glucose (Sigma) with 15% heat inactivated dialysed FCS (Thermofisher). Subsequently, FLDCS were resuspended in the same media + addition of defined: glucose (Agilent) fructose, dextrin, inosine, glycerol-3-phosphate, Mannose or galactose (all ThermoFisher) concentrations. FLDCs were pre-incubated for 2h at 37 °C, 5% CO_2_ followed by the addition of either swollen arrested Af spores at an MOI of 5:1 or equal volume media only controls.

In FLDC omnilog experiments, cultured FLDCs were washed twice in IF-M1 (IF-M1, Biolog, 5% dialysed FCS, 0.2 mM L-Glutamine) media followed by plating on a Phenotype MicroArray Mammalian plate PM-M1 (Biolog) at 1 × 10^5^ cells/well in IF-M1 media. Following a 2h pre-incubation at 37 °C at 5% CO_2_, either PFA-Killed swollen *Af* spores (resuspended in the Biolog Redox Dye (Biolog)) at an MOI of 5:1 or redox dye only controls were added to appropriate wells of FLDC PM-M1 plates and subsequently incubated at 37 °C + 5% CO_2_ for 12 h. Following incubation, OD_590_ and OD_750_ measurements were taken utilising a TECAN Spark multimode microplate reader (Tecan). To ascertain ability of individual nutrient wells to support FLDC metabolism, OD_750_ measurements were first subtracted from OD_590_ readings. The average reading from the 3 negative control wells were then subtracted from well readings and the percentage of the average reading from the 3 α-D-Glucose (the nutrient supporting the most FLDC metabolism) wells calculated and plotted.

### Metabolic profiling

SCENITH™ reagents kit (DG, O, puromycin and anti-puromycin AF488 antibodies) were a kind gift from (R.A). For *in vivo* and FLDC single cell metabolic profiling of cell, SCENITH was performed as previously described^57^. Briefly, lung (in vivo) or FLDC single cell suspensions were split equally into 4/ sample and treated with Control (DMSO), 2-Deoxy-Glucose (2-DG; 100DmM), Oligomycin (O; 1 µM) for 15 mins or a combination of 2DG (5 minutes) followed by Oligomycin (DGO) ( a further 10 minutes) Following metabolic inhibitor treatment periods, Puromycin (final concentration 10 µg/mL) was added to cultures for a further 30 Dmin. After puromycin treatment, cells were washed in ice-cold lung buffer and stained in Fc-Block (BD Biosciences) and Live Dead Blue fixable viability dye (ThermoFisher) for 15Dminutes at 4 °C in Lung buffer. Subsequently, extracellular surface markers were stained for 45 min at 4 °C before cells were fixed in 1% PFA (Sigma) for a further 10mins. Samples were then fixed/permeabilised using Foxp3/transcription factor Buffer Set (ThermoFisher) for 10mins, blocked in permeabilisation buffer + 10% FCS (both Sigma) for 15 mins, followed by intracellular staining of puromycin for 1 h in diluted (10×) permeabilization buffer at 4 °C. Finally, samples were acquired using the Cytek Aurora flow cytometer. gMFIs of puromycin served as translation rate for Co, DG, O and DGO conditions and utilised to calculate: metabolic activity (C-DGO), Glucose dependency ((100(Co-DG)/(Co-DGO)) and mitochondrial dependency ((100(Co-O)/(Co-DGO)), glycolytic capacity (100-((100(Co-O)/(Co-DGO)) and FA oxidation (FAO) capacity (100-((100(Co-O)/(Co-DGO)) and glycolytic capacity.

Extracellular flux analysis was performed with the Seahorse XFe-96 Flux Analyzer (Agilent) to measure oxygen consumption (OCR) and extracellular acidification rates (ECAR) of FLDCs. FLDCs were harvested in RPMI1640 Glutamax + 10% FCS, washed in RPMI-1640 only (sigma) and glucose starved by loading 2x10^5^ FLDC/well in RPMI-1640 for 2 hours. Following glucose starvation, media was changed for XF media (RPMI-1640, 1 mM HEPES without bicarbonate buffer, 1 mM pyruvate, 2 mM L-Glutamine, pH 7.4. All Agilent) and degassed at 37 °C and atmospheric levels of CO_2_ for 1 hour. After 4 baseline measurement cycles on the Seahorse, 1 million killed spores (MOI 5) were injected, and 14 further measurement cycles were performed.

### Flow cytometry and Sorting

Surface and intracellular cytokine staining were undertaken as previously described^11^. Briefly, immune cell cytokine secretion was ascertained by stimulating lung single cell suspensions with PMA (30 ng/ml, Sigma), Ionomycin (1 μg/ml, Sigma) and GolgiStop (BD) for 3 hours prior to surface and intracellular staining. Surface staining was undertaken on equal numbers of cells per sample. Following a wash with ice-cold PBS, viability was ascertained by incubation in Live/Dead Blue (ThermoFisher) for 15 min at 4C and subsequently incubated in Fc-Block (BD Biosciences) and surface markers (antibodies and chemokines listed in supplementary table 1 in FACS buffer (PBS containing 2% FBS and 2 mM EDTA, all sigma) for 1 hour at 4C. Post staining, cells were washed and fixed in 1% PFA in PBS (both sigma) for 10 min at RT. For detection of intracellular antigens, extracellular labelled cells were processed with Foxp3 buffer staining set as per manufacturer’s protocol (ThermoFisher) and stained with selected antibodies. Samples were acquired either on a Cytek Aurora flow cytometer or Attune Nxt Acoustic focusing cytometer (ThermoFisher). CountBright Absolute Counting Beads (ThermoFisher) were added to samples before acquisition to accurately enumerate cell populations.

In experiments where lung dendritic cells (DCs) were sorted prior to RNA-sequencing, lung single cell suspensions were generated by pooling 3-4: PBS, 6x dosed or 9x dosed mice per sample. Samples were first enriched on an Optiprep density gradient (ThermoFisher): single cell suspensions were resuspended in 1.085g/mL Optiprep and subsequently layered with 1.065 g/mL Optiprep and HBSS (Sigma). Cells were centrifuged at 600xg for 15 minutes and enriched DCs extracted from the cell layer formed between the 1.065 g/ml and HBSS layers. Cells were washed in FACS buffer, extracellular surface markers stained (as described above) and resuspended in ice-cold FACS buffer for onward sorting utilising the Bigfoot Spectral Cell sorter (ThermoFisher) in purity mode with the 100μm nozzle. 100 Spore^+^ and Spore^-^ Mgl2^+^CD209a^+^ cDC2s were sorted directly into 5ul 1x NEBnext Cell lysis buffer (New England Biolabs), snap frozen on dry ice and stored at -80C prior to onward RNAseq library amplification and preparation.

### RNAseq and analysis

Amplification and subsequent library preparation from the 100 Spore^+/-^ Mgl2^+^CD209a^+^ cDC2s sorted into 1x NEBnext Cell lysis buffer (New England Biolabs) was undertaken utilising NEBNext® Single Cell/Low Input RNA Library Prep Kit for Illumina® (New England Biolabs) as per manufacturer’s instructions. Prepared samples were subsequently sequenced on the Illumina NovaSeq 6000 Sequencing System utilising a configuration of HiSeq 2x150bp HiSeq 2x150b. For analysis, Sequence reads were trimmed to remove possible adapter sequences and nucleotides with poor quality using Trimmomatic v.0.36. The trimmed reads were mapped to the Mus musculus GRCm38 reference genome available on ENSEMBL using the STAR aligner v.2.5.2b. Unique gene hit counts were then calculated by using featureCounts from the Subread package v.1.5.2. The hit counts were summarised and reported using the gene_id feature in the annotation file. Only unique reads that fell within exon regions were counted. Using the generated hit counts table, differential gene expression analysis was undertaken using DESeq2. The Wald test was used to generate p-values and log2 fold changes. Genes with an adjusted p-value < 0.05 and absolute log2 fold change > 1 were called as differentially expressed genes for each comparison. Gene enrichment analysis was performed with the clusterProfiler package for R, using the ‘compareCluster’ function with fun = “gseKEGG” and pvalueCutoff = 0.05. Pathways were considered statistically significant where adjusted p value <0.05. Pathways associated with DC function were selected for display in heatmaps.

### Metabolomics and Airway glucose measurement

PBS and 9x *Af* dosed mice from the anti-fungal allergic inflammation model described above were sacrificed, followed by immediate BAL with 1ml of ice-cold PBS (Sigma). Samples were flash frozen on dry ice and stored at -80°C until processing for Liquid Chromatography-Mass Spectrometry (LC-MS), with all utilised solvents Fisher Optima LC-MS grade (Fisher Scientific, UK). For airway glucose measurement collected BAL samples were thawed and glucose concentration ascertained utilising an AmplexRed Glucose/Glucose oxidase assay kit (Thermofisher) as per manufacturers protocol.

For LC-MS metabolomics we modified protocols as previously described^109^, briefly samples were thawed on ice and quenched, protein removed (crash), and more polar metabolites extracted into the supernatant by addition of ultra cold LC-MS grade methanol followed by vortexing for 1 min. Samples were next centrifuged at 17000 *g* for 15 min at 4°C using a Thermo Scientific Fresco Heraeus 17 centrifuge (Thermo Electron LED GmbH, Germany) to pellet and remove solid content with extracted supernatant dried to completion overnight in a Scanvac Maxi system (Labogene A/S, Denmark). Where LC-MS was not immediately undertaken, samples were stored as dried extract at -80°C. Blank samples consisting of LC-MS grade water were treated equivalently.

Dried extracted samples were reconstituted in 60 µL of LC-MS grade water and centrifuged at 17000 x*g* for 15 min at 4°C .40 µL of sample supernatant were transferred to glass vials fitted with 300 µL fused inserts (Chromatography Direct VI-04-12-02RVAE; Chromatography Direct Ltd., Runcorn, U.K.), with the remaining liquid content pooled across samples to generate a stock biological pool sample for use as: column conditioning, biological QCs and ddMS^2^ samples within the analysis acquisition sample set, as described121121. All samples, blanks, process blanks and QCs were loaded into sample racks and placed into an autosampler (set to 4°C) of a Thermo Vanquish UHPLC system (Thermo Scientific, DionexSoftron GmbH, Germany) prior to data acquisition.

Reversed phase data acquisition for all samples was undertaken on a Thermo Vanquish UHPLC system in conjunction with a Thermo Orbitrap IDX mass spectrometer (ThermoFisher Scientific, U.S.A.). The UHPLC method utilised is a standard CMR protocol as previously described122122. Briefly, the UHPLC method involved a 5 µL sample injection into a 0.4 µL/min flow gradient of mobile phase A (water + 0.1% formic acid) and mobile phase B (methanol + 0.1% formic acid). The column used was a Thermo Hypersil Gold aQ 100x2.1 mm, 1.9 µm column set at 50°C. LC-MS data were first acquired for all samples in positive electrospray (ESI) ionisation mode (ESI+) then for all samples in negative ESI ionisation mode (ESI-). The order of sample injection was block randomised and QCs were run following every fifth sample injection. The normal MS acquisitions for each sample were of type MS^1^, utilising an Orbitrap resolution of 120,000 and scan range *m/z* 66.7 to 1000. ddMS^2^ acquisitions of pool samples were run as a set of four injections once all samples (in each ionisation mode) had been injected. The ddMS^2^ acquisitions utilised a “precursor” Orbitrap resolution of 60,000, a fragment ion resolution of 30,000 and in-method stepped HCD Collision Energies of 20, 40 and 60. Data were acquired for four precursor ion mass ranges (one range per injection); specifically at *m/z* 66.7-1000, 66.7-300, 300-600 and 600-1000.

In each ion mode the standards were run as a group of injections either side of the sample injections – not intermingled with the samples – plus separated from the samples by several column conditioning QC samples. Aside from using injection volumes of 1 µL, the acquisition methods for the standards were identical to those used for the samples; with ddMS^2^ only being undertaken at the end of the standard group following the sample injections.

Following data acquisition, raw data were deconvoluted utilising compound discoverer 3.3 software (Thermal) and a peak table generated for each set of data. The peak table was then imported into MATLAB 2023a (MATHWORKS, MA) for multivariate analysis. QC correction was applied to the data as previously described^109^. Data was then log-scaled and subjected to principal component analysis (PCA). Partial least squares for discriminant analysis (PLS-DA) were also applied to assess the statistical significance of the separation between groups. The PLS-DA models were validated by using a bootstrapping couple with permutation tests method as previously described^112^.

### Statistical analysis

Statistical analysis was performed using Prism v9 (GraphPad). Data are shown as mean valuesD±DSEM. Where applicable, data were analysed by unpaired t-test, paired t-test or one-way analysis of variance (ANOVA) with Tukey’s post-test as appropriate. Significant differences were defined at PD<0.05. Statistical analysis of RNA-seq and metabolomics data is detailed in appropriate sections above.

## Data Availability

The data that support the findings of this study are available from the corresponding author upon request. RNA-seq data is currently being deposited at Gene Expression Omnibus. Metabolomics data is currently being deposited at EBI’s MetaboLights.

## Acknowledgements

We thank members of the MRC Centre for Medical Mycology at the University of Exeter for scientific discussions and some experimental assistance. Additionally, we thank the University of Exeter Centre for Cytomics and University of Exeter Biological Services Unit for technical and experimental assistance. P.C.C. and J.F-S. were supported by funding from a Wellcome Trust Sir Henry Dale Fellowship (218550/Z/19/Z), the MRC Centre for Medical Mycology at the University of Exeter (MR/N006364/2 and MR/V033417/1), and the NIHR Exeter Biomedical Research Centre (NIHR203320). P.T.N and D.C. were supported by an MRC Doctoral Training Grant MR/P501955/2. The views expressed are those of the author(s) and not necessarily those of the NIHR or the Department of Health and Social Care. Schematics in figures were created in https://BioRender.com. For the purpose of open access, the author has applied a ‘Creative Commons Attribution (CC BY) licence to any Author Accepted Manuscript version arising from this submission’.

## Author contributions

P.C.C. was responsible for conceptualisation. J.F-S., G.V., P.T.N, N.G., Y.X., S.A.P.C, D.C., B.L.M. performed the experiments or undertook data analysis. R.A., S.A., S.O., S.L-L, R.G., D.A-J, E.B., G.B. provided resources J.F-S, G.V., P.C.C. prepared the manuscript with input from all authors. P.C.C. obtained funding for study.

