## Supplemental Figures and Table for "Mgl2^+^ cDC2 triggering of fungal allergic inflammation depends on a spore induced glycolytic shift fuelled by local availability of glucose"

**Supplementary Methods**

**ELISAs**

Wells of Nunc 96-well plates (Thermofisher) were coated with capture antibody and incubated overnight at 4$^{o}$C. Plates were washed three times (PBS + 0.05% Tween-20 (Sigma)), blocked with PBS + 10% newborn calf serum (NCS) (Sigma) for 90 mins at 37$^{o}$C followed by the addition of relevant samples. Following overnight incubation at 4$^{o}$C, plates were washed and biotin-conjugated detection antibody added for 1 hour at 37$^{o}$C with the subsequent addition streptavidin-horseradish peroxidase (Seracare) (diluted 1000-fold in PBS + 1% NCS) for 30 mins at 37$^{o}$C. After thorough washing, 0.66 mM 3,3',5,5'-Tetramethylbenzidine (Sigma) dissolved in phosphate-citrate buffer (0.5 M, pH 5.0) was added and allowed to develop for maximal signal. The reaction was then stopped with 0.18 M H2SO4, followed by reading of fluorescence on a Tecan plate reader at 450 nm with background 570 nm fluorescence subtracted. List of antibodies utilised are listed in Supplementary Table 2.

**Supplementary figure legends:**

**Supplementary Figure 1. Spore uptake is dominated by airway CD45$^+$ immune cells.**
Mice were exposed to labelled *A. fumigatus* (*Af*) spores (strain: CEA10, 4 x $10^5$ spores) via intranasal (i.n.) transfer, 3h later following CD45 administration i.n., lung tissue was harvested for Precision Cut Lung Slices (PCLS). **A**) Analysis pipeline to quantify spore (white) uptake by CD45$^+$ cells (purple), EPCAM$^+$/Podoplanin$^+$ epithelial cells (green) and extracellular spores. Whole mapping of PCLSs were performed utilising confocal microscopy at 10x magnification. 6-8 fields of view (FOVs), representative of the whole lung slice, were then assigned and 20x magnification z-stacks acquired. Spore and cell surfaces were rendered using IMARIS software to facilitate quantitative analysis. **B**) Representative PCLS renderings demonstrating epithelial *Af* spore uptake (left) and extracellular spores (right).

**Supplementary Figure 2. Repeated exposure to spores induces hallmarks of fungal allergic inflammation.**
Naïve mice or mice that were repeatedly exposed to unlabelled *Af* via intranasal transfer on the indicated time points, followed by exposure to GFP-expressing *Af* on the final dose. Tissues were harvested 3h after the first, sixth or ninth dose of spores (1x, 6x and 9x, respectively). **A**) Gating strategy utilised to identify lung immune cell subsets from flow cytometric analysis **B**) Quantification of lung immune subsets. Graphs display the total numbers and percentages of CD45$^+$ cells for alveolar macrophages (AlvM), interstitial macrophages (IntM), DCs, monocytes, eosinophils and neutrophils from the lung tissue. **C**) Gating strategy utilised to identify cytokine-producing lung CD4+ T cells. **D**) Quantification of lung CD4$^+$ T-cell cytokine expressing cell. Graphs display cytokine$^+$ T-cells as as a percentage of total lung CD4$^+$ T cells.
**B** - **D**) data are from two independent experiments (n = 4 - 8 biologically independent animals per group).
Significance was determined by 1-way ANOVA with Tukey's multiple-comparison post-hoc test between indicated groups *P < 0.05, **P<0.01, **** P<0.0001, NS not significant.

**Supplementary Figure 3. Unbiased clustering analysis to define spore uptake by lung immune cell populations following *Af* exposure.**
FlowSOM and tSNE analysis was undertaken on *Af* spore$^+$ lung CD45$^+$ cells, FlowSOM metaclusters were merged and identified based on marker expression to resolve immune subsets. tSNE plots show marker expression of the concatenated FCS files with representative examples shown. Tissues were harvested 3h after the first, sixth or ninth dose of spores (1x, 6x and 9x, respectively).

**Supplementary Figure 4. Repeated exposure to spores reduced AlvM, and increased DC and granulocyte, *Af* uptake.**
Naïve mice or mice that were repeatedly exposed to unlabelled *Af* via intranasal transfer on the indicated time points, followed by exposure to GFP-expressing *Af* on the final dose. Tissues were harvested 3h after the first, sixth or ninth dose of spores (1x, 6x and 9x, respectively). **A**) Total number of spore$^+$ lung from defined exposures. **B**) Graphs display spore uptake by lung immune cell populations plotted as percentage of spore$^+$ cells(top) or percentage of each immune subset that were spore$^+$ (bottom).
**B**) data are from two independent experiments (n = 6 - 8 biologically independent animals per group).
Significance was determined by 1-way ANOVA with Tukey's multiple-comparison post-hoc test between indicated groups *P < 0.05, **P<0.01, ***P<0.001, **** P<0.0001, NS not significant,

**Supplementary Figure 5. Unbiased clustering analysis to define spore uptake by lung DC subsets following *Af* exposure.**

FlowSOM and tSNE analysis was undertaken on *Af* spore$^{+}$ lung DCs (Lin$^{-}$Mertk$^{-}$CD64$^{-}$SiglecF$^{-}$ Ly6G$^{-}$CD11c$^{+}$MHCII$^{+}$). FlowSOM metaclusters were merged and identified based on marker expression to resolve individual DC subsets. tSNE plots show marker expression of the concatenated FCS files, with representative examples shown. Tissues were harvested 3h after the first, sixth or ninth dose of spores (1x, 6x and 9x, respectively).

**Supplementary Figure 6. Mgl2$^{+}$ cDC2s are the dominant DC subset to acquire spores following *Af* exposure.**

Naïve mice or mice that were repeatedly exposed to unlabelled *Af* via intranasal transfer on the indicated time points, followed by exposure to GFP-expressing *Af* on the final dose. Tissues were harvested 3h after the first, sixth or ninth dose of spores (1x, 6x and 9x, respectively). **A**) Graphs display the total number of cells in each DC subsets (top) or spore uptake plotted as percentages of whole lung spore$^{+}$ population, percentages of all spore$^{+}$ DCs events and percentages of each DC subset that were spore positive. **B**) Quantified CD86 and CD40 expression of defined DC subsets. Graphs show the fold change in expression of spore$^{+}$ versus spore$^{-}$ DCs of each DC subset lines indicate relevant DC populations from the same mouse.

**A** – **B**) data are from two independent experiments (n = 6 - 8 biologically independent animals per group).

Significance was determined by **A**) 1-way ANOVA with Tukey's multiple-comparison post-hoc test or **B**) a paired Student's t test between indicated groups *P < 0.05, **P<0.01, **** P<0.0001, NS not significant.

**Supplementary Figure 7. Spore acquisition by Mgl2$^{+}$CD209a$^{+}$ cDC2s drives an increased metabolic transcriptional profile.**

Mice were repeatedly exposed to unlabelled *Af* spores (4 x $10^{5}$ per dose) via intranasal for 5 or 8 doses followed by a final dose of labelled *Af*. An additional control group of mice received 9 doses of PBS only. Tissues were harvested 3h post sixth and ninth dose of spores (6x and 9x, respectively). Spore$^{-}$ and spore$^{+}$ Mgl2$^{+}$CD209a$^{+}$ cDC2s were isolated from the lung tissue of mice that received 6x and 9x doses of *Af*, alongside Mgl2$^{+}$CD209a$^{+}$ cDC2s from the lung tissue of PBS exposed mice. Transcriptome of each population was generated using low input kits followed by RNA sequencing (Illumina). KEGG pathway analysis show significant up or down-regulated transcriptional pathways of indicated spore$^{+}$ Mgl2$^{+}$CD209a$^{+}$ cDC2s from *Af*-exposed mice versus spore$^{-}$ Mgl2$^{+}$CD209a$^{+}$ cDC2s from *Af*-exposed mice. Circles indicate p.adjust < 0.05 compared to spore- counterparts calculated via Wald test in DESeq2.

Each individual sample is a pool from 3-4 biologically independent animals derived for 3 independent experiments.

**Supplementary Figure 8. Spore uptake impacts Mgl2$^{+}$CD209a$^{+}$ cDC2s transcript expression, especially metabolic pathways.**

Mice were repeatedly exposed to unlabelled *Af* spores (4 x $10^{5}$ per dose) via intranasal for 5 or 8 doses followed by a final dose of labelled *Af*. An additional control group of mice received 9 doses of PBS only. Tissues were harvested 3h post sixth and ninth dose of spores (6x and 9x, respectively). Spore$^{-}$ and spore$^{+}$ Mgl2$^{+}$CD209a+ cDC2s were isolated from the lung tissue of mice that received 6x and 9x doses of *Af*, alongside Mgl2$^{+}$CD209a$^{+}$ cDC2s from the lung tissue of PBS exposed mice. Transcriptome of each population was generated using low input kits followed by RNA sequencing (Illumina).Heatmaps show scaled gene expression of KEGG term enriched genes associated with: **A**) DC function, **B**) Signalling pathways, **C**) Metabolic pathways from or **D**) Curated gene lists from associated metabolic pathways of indicated spore$^{+/-}$ Mgl2$^{+}$CD209a$^{+}$ cDC2s from *Af*-exposed mice versus Mgl2$^{+}$CD209a$^{+}$ cDC2s from PBS

mice. Circles indicate p.adjust < 0.05 compared to PBS controls calculated via Wald test in DESeq2.
Each individual sample is a pool from 3-4 biologically independent animals derived for 3 independent experiments.

**Supplementary Figure 9. Spores and zymosan exposure trigger activation of FLDC subsets and secretion of pro-inflammatory factors.**
Flt3L bone marrow derived DCs (FLDCs) generated from mice were exposed to labelled 4-hour swollen Δ*pyrG Af* spores or zymosan. **A**) Gating strategy for the identification of FL-DC subsets (FL-pDC, FL-cDC1, and FL-cDC2) and identification of spore$^{+}$ FL-cDC2s compared to untreated FL-cDC2s (media). **B**) Representative flow cytometry overlay plots and graphs show the gMFI of co-stimulatory molecules (CD40 and CD86), CCR7 and MHCII of each FLDC subset following 6h incubation with *Af* spores or zymosan. **C**) Representative flow cytometry overlay plots and graphs show the gMFI of co-stimulatory molecules (CD40 and CD86), CCR7 and MHCII of FL-cDC2s following 2h or 6h incubation with *Af* spores or zymosan. **D**) FLDC chemokine and cytokine secretion following 2h or 6h incubation with *Af* spores or zymosan.
**B - C**) Data are from three independent experiments (CD40, CD86, MHCII, n = 10-15 FLDCs derived from biologically independent animals per group) or one independent experiment (CCR7, n = 4 FLDC cultures from biologically independent animals per group)
**D**) Data are from two independent experiments (n = 5-8 FLDCs derived from biologically independent animals per group)
Significance was determined by 1-way ANOVA with Tukey's multiple-comparison post-hoc test between indicated groups *P < 0.05, **P<0.01, ***P<0.001, **** P<0.0001, NS not significant.

**Supplementary Figure 10. FLDC metabolic activity rapidly increases following exposure to spores.**
**A**) Δ*pyrG Af* spores were swollen for 4 hours (arrested swollen live) and some killed with 1% PFA (1% PFA-killed Af). Both groups were injected into Agilent SEAHORSE XF analysers to track ExtraCellular Acidification Rate (ECAR) and Oxygen Consumption Rate (OCR) tracked over time. **B**) Flt3L bone marrow derived DCs (FLDCs) generated from mice were exposed to 4h arrested swollen live *Af* or 4h 1% PFA-killed *Af*. Representative flow cytometry overlay plots and graphs show the FL-cDC2 gMFI of co-stimulatory molecules (CD40 and CD86), CCR7 and MHCII following 6h incubation. **C)** FLDCs were ran on the Agilent SEAHORSE XF analysers and exposed to acute injections of media, swollen Af or zymosan. Left graph shows normalised ECAR and OCR of FLDCs over time while right graph shows the change in ECAR and OCR between the pre-injection measurement cycle and the final cycle.
**A**) Data are from one of two independent experiments (n = 3 wells per group)
**B**) Data are from one of two independent experiments (n = 3 FLDCs derived from biologically independent animals per group)
**C**) Data are from two independent experiments (n = 20 wells per group, each experiment utilised FLDCs pooled from two biologically independent animals).
Significance was determined by 1-way ANOVA with Tukey's multiple-comparison post-hoc test between indicated groups. **** P<0.0001.

**Supplementary Figure 11. FL-cDC2s are the dominant FLDC subset to acquire spores.**
Flt3L bone marrow derived DCs (FLDCs) generated from mice were exposed to 4-hour swollen Δ*pyrG Af* spores for 2 or 6 hours. Representative flow plots (**A**) and graph (B) shows spore uptake by FL-cDC1, FL-cDC2 and FL-pDC subsets.
**A - B**) Data are from two independent experiments (N=9 FLDCs derived from biologically independent mice per group)

Significance was determined by 1-way ANOVA with Tukey's multiple-comparison post-hoc test between indicated groups. **** P<0.0001.

**Supplementary Figure 12. Inhibition of glycolysis reduces FLDC secretion of cytokines but not growth of spores.**

**A**) Flt3L bone marrow derived DCs (FLDCs) generated from mice were exposed to labelled 4-hour swollen Δ*pyrG Af* spores in the presence or absence of various metabolic inhibitors. Graph shows FLDC cytokine secretion after 6h incubation. **B**) *Af* spores were germinated in PBS or 2-DG for 20 hours. Graphs show percentage of spores germinated measured by flow cytometry (left) and metabolic activity measured by addition of resazurin to cultures.

**A**) Data are from two independent experiments (n = 8 FLDCs derived from biologically independent mice per group)

**B**) Data are from one independent experiment (n = 4 wells of *Af* spores per group)

Significance was determined by 1-way ANOVA with Tukey's multiple-comparison post-hoc test between indicated groups *P < 0.05, **P<0.01, ***P<0.001, **** P<0.0001.

**Supplementary Figure 13: FL-cDC2 *Af* spore responses are dependent on CLR signalling.**

**A**) Flt3L bone marrow derived DCs (FLDCs) generated from WT mice were exposed to media to labelled 4-hour swollen Δ*pyrG Af* spores in the presence of R406 Syk inhibitor or DMSO vehicle control for 12 hours. Graphs display cDC2 activation marker (CD86 and CD40) and MHCII expression normalised as fold change versus media only exposed controls. **B)** FLDCs generated from WT and various C-type lectin receptor knockout mice were exposed to spores (bottom) for 12 hours. Graphs display cDC2 activation marker (CD86, CD40 and CD80) expression normalised as fold change from respective media only exposed controls.

**A**) Data are from two independent experiments (n = 8 FLDCs derived from biologically independent mice per group)

**B**) Data are from 3 independent experiments (n = 3-15 FLDCs derived from biologically independent mice per group)

Significance was determined by 1-way ANOVA with Tukey's multiple-comparison post-hoc test between indicated groups *P < 0.05, **P<0.01, ***P<0.001, **** P<0.0001.

**Supplementary Figure 14. Repeated exposure to spores increases glycolysis of lung myeloid and granulocyte cell populations.**

Mice were repeatedly exposed to PBS or *Af* spores (4 x $10^5$ per dose) via intranasal for 9 doses (9x *Af*). Tissues were harvested 24h post ninth dose of spores with whole lung suspensions processed for SCENITH metabolic analysis via flow cytometry. Each sample is independently treated ex vivo with control (Co), 2-DG, Oligomycin-A or 2-DG + Oligomycin, gMFI of puromycin staining (rate of translation) of each treatment indicates metabolic activity, glucose dependency, glycolytic capacity, mitochondrial dependency and fatty acid oxidation (FAO) capacity). **A**) Representative flow cytometry overlay plots of gMFI puromycin staining of lung interstitial macrophages (IntM), monocytes, neutrophil and eosinophil populations from PBS and 9x *Af* dosed animals.

**B**) Graphs displaying metabolic activity (measured as Co-DGO Puromycin gMFI) and **C)** Corresponding metabolic dependency and capacity parameters of PBS and 9x *Af* dosed animals.

**B - C**) Data are from one independent experiment (n = 3 biologically independent mice per group).

Significance was determined by 1-way ANOVA with Tukey's multiple-comparison post-hoc test between indicated groups *P < 0.05, **P<0.01, ***P<0.001, **** P<0.0001.

**Supplementary Figure 15. Uptake of spores by FL-cDC2s depend on glucose, but not other carbon sources.**

**A**) Flt3L bone marrow derived DCs (FLDCs) generated from WT mice were precultured in basal media prior to addition of labelled 4-hour swollen Δ*pyrG Af* spores in the presence of specified nutrients. **B**) Graphs display FL-cDC2 viability and spore uptake after exposure to spores whilst individually supplemented with glucose, fructose, dextrin, inosine, glycerol-3-phosphate, mannose or galactose after 6h incubation.

**B**) Data are from two independent experiment (n = 7-8 FLDCs derived from biologically independent mice per group)

Significance was determined by 1-way ANOVA with Tukey's multiple-comparison post-hoc test between indicated groups **P<0.01, ***P<0.001, **** P<0.0001.

**Supplementary Figure 16. Availability of glucose does not impact FLDC cDC2 viability or uptake of spores.**

**A**) Flt3L bone marrow derived DCs (FLDCs) generated from WT mice were precultured in basal media prior to addition of labelled 4-hour swollen Δ*pyrG Af* spores in the presence of different concentrations of glucose. **B**) Graphs display FL-cDC2 viability and spore uptake after exposure to spores in different concentrations of glucose after 6h incubation.

**B**) Data are from two independent experiments (n = 7-8 FLDCs derived from biologically independent mice per group)

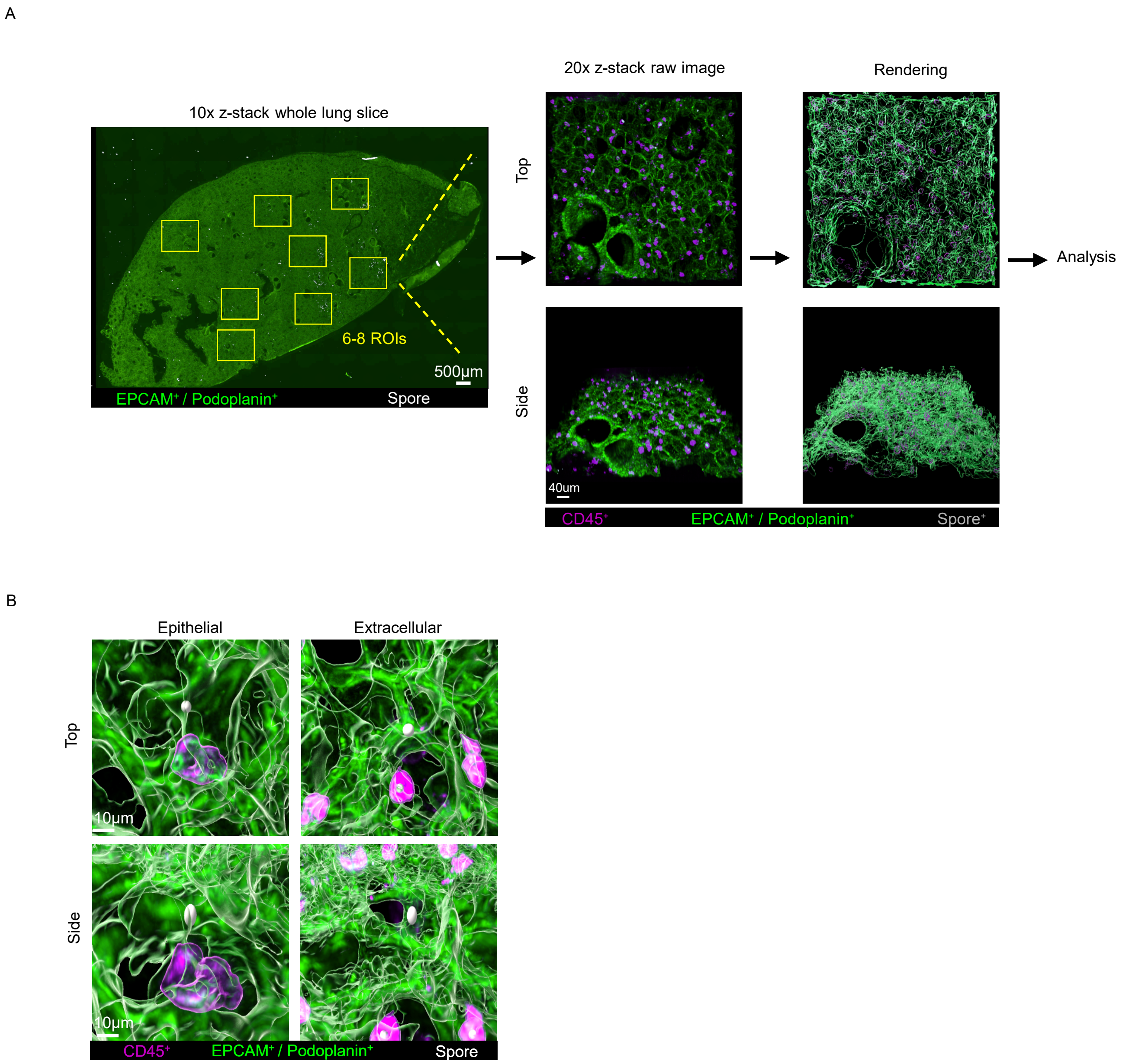


Supp. Figure 1

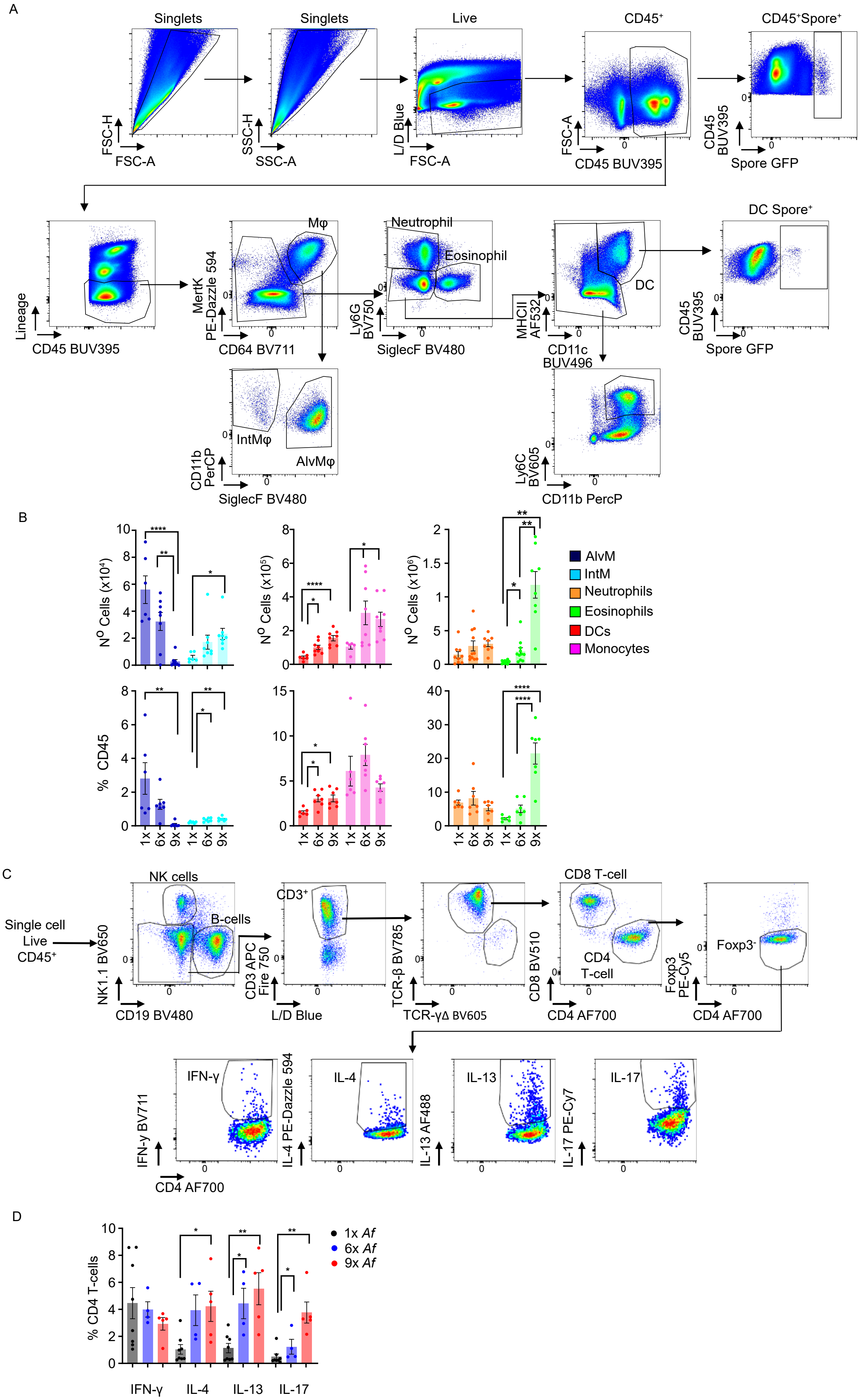


Supp. Figure 2

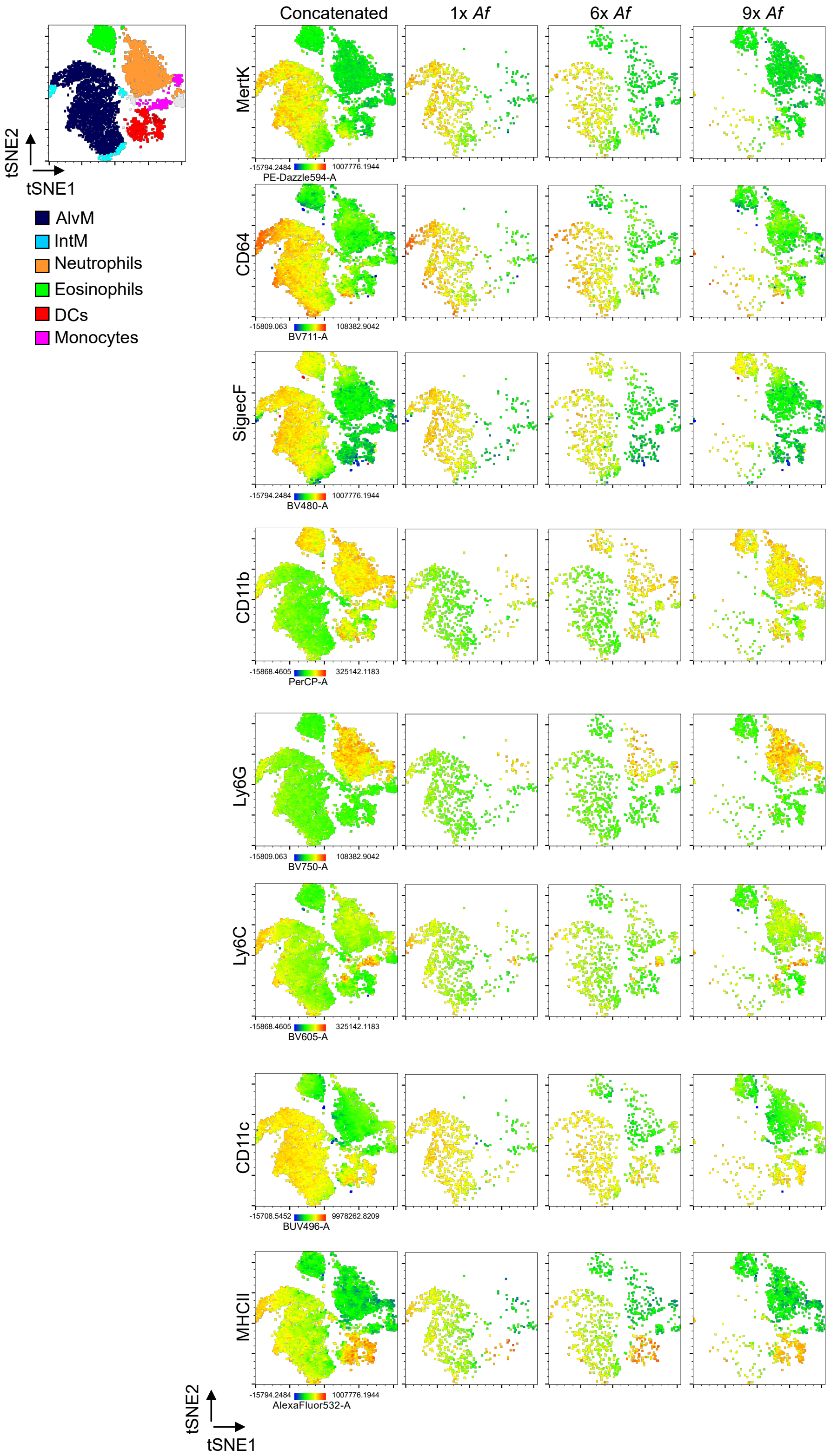


Supp. Figure 3

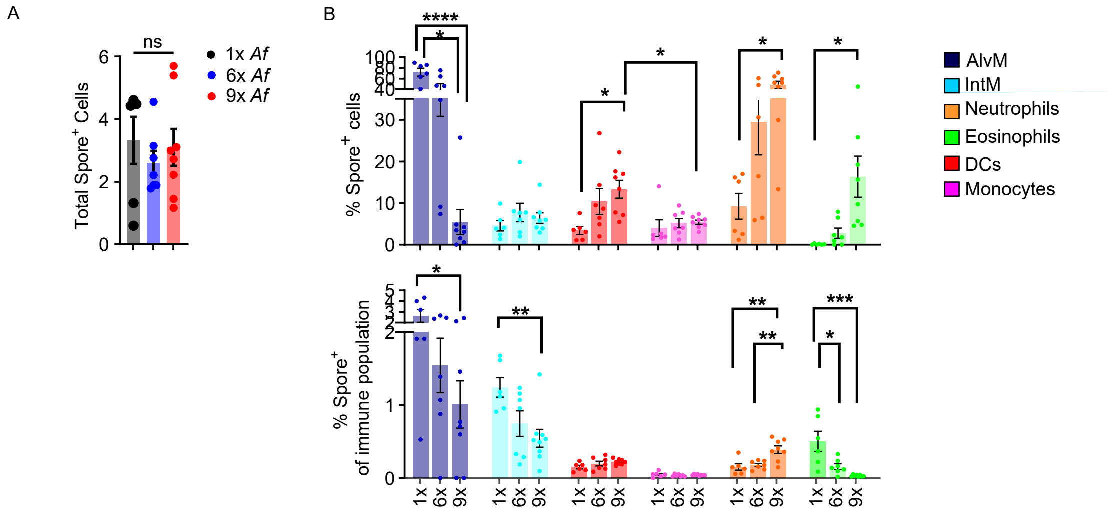


Supp. Figure 4

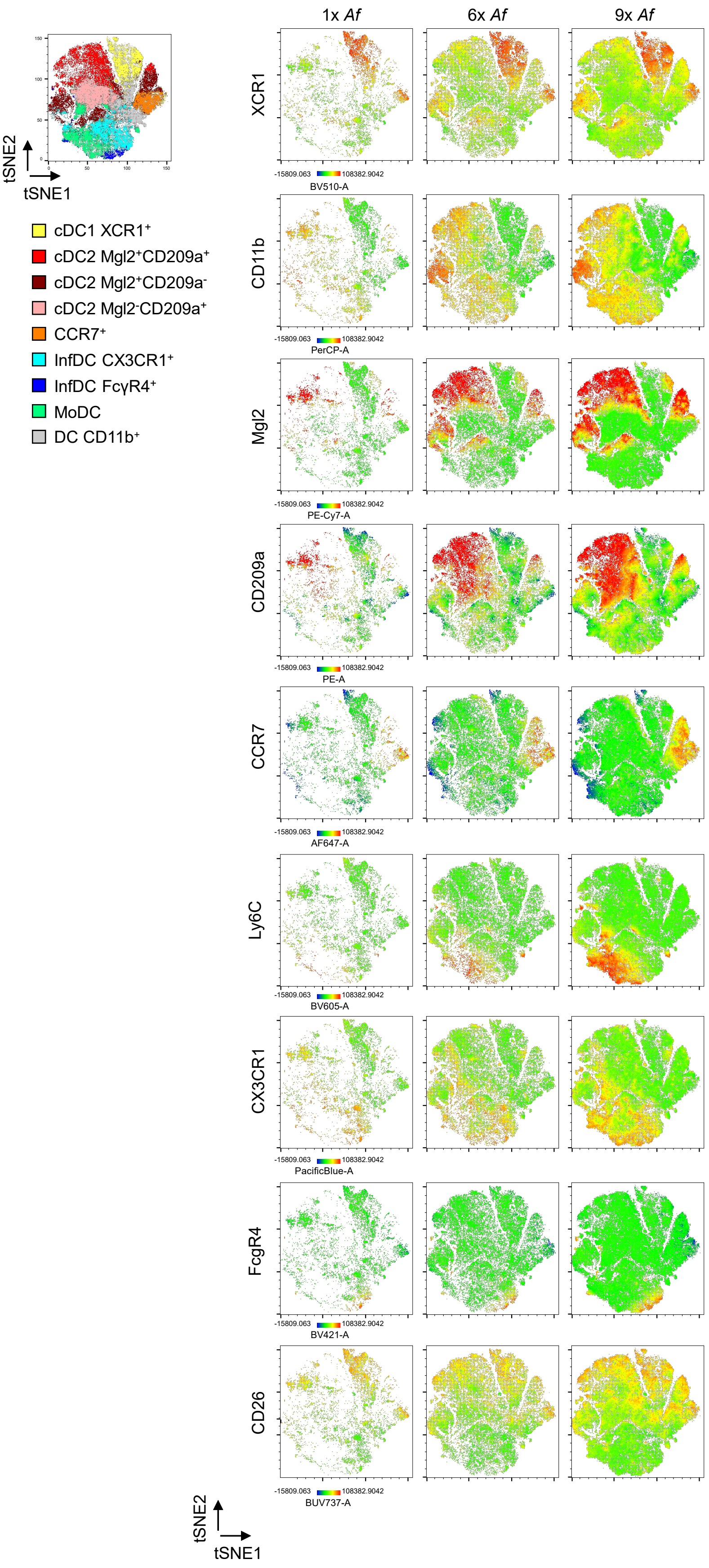


Supp. Figure 5

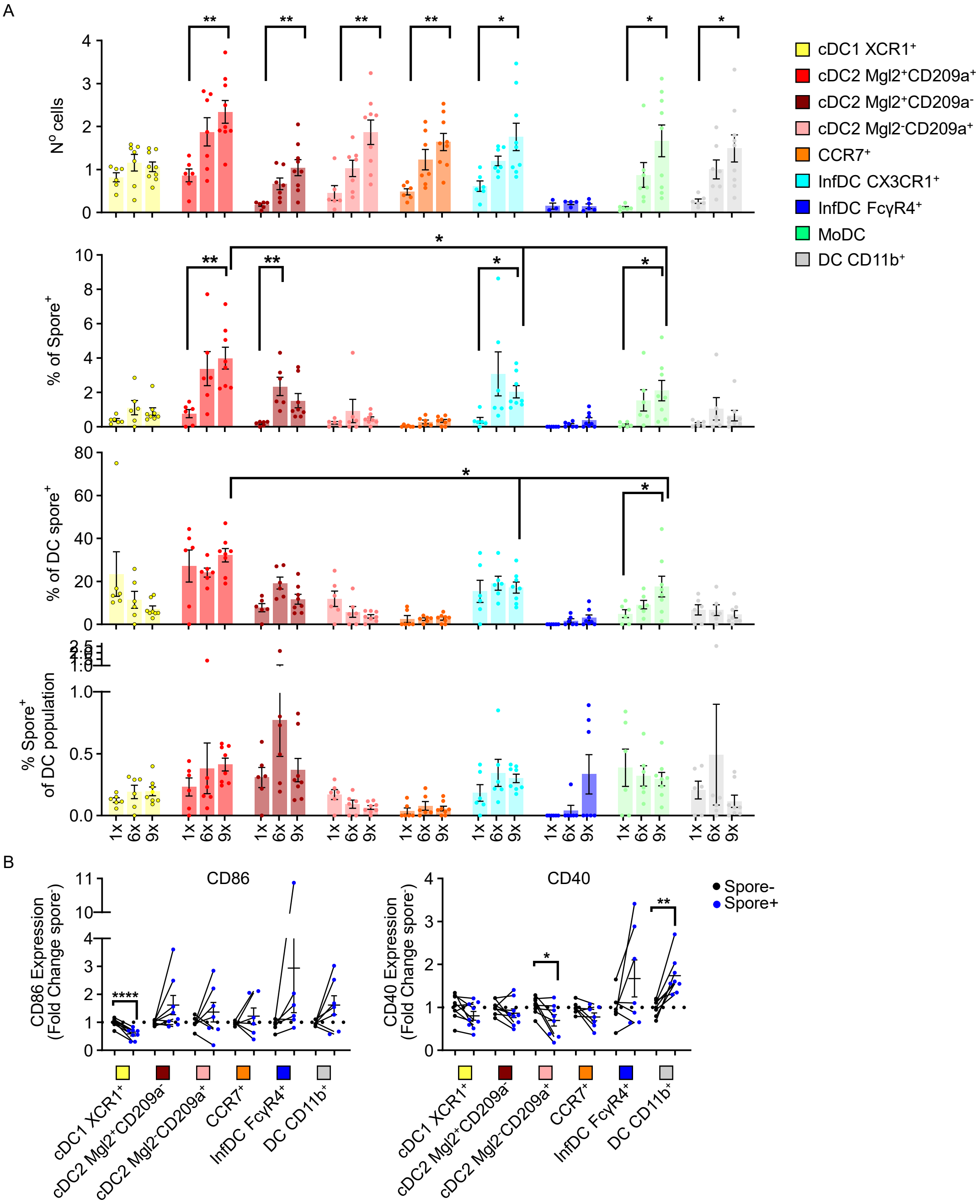


Supp. Figure 6

A

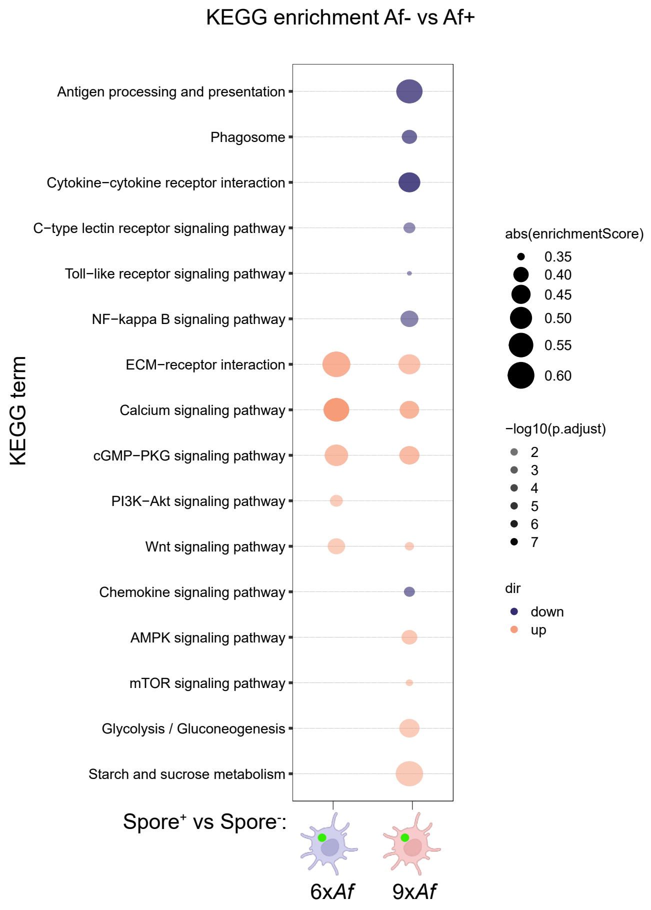


Supp. Figure 7

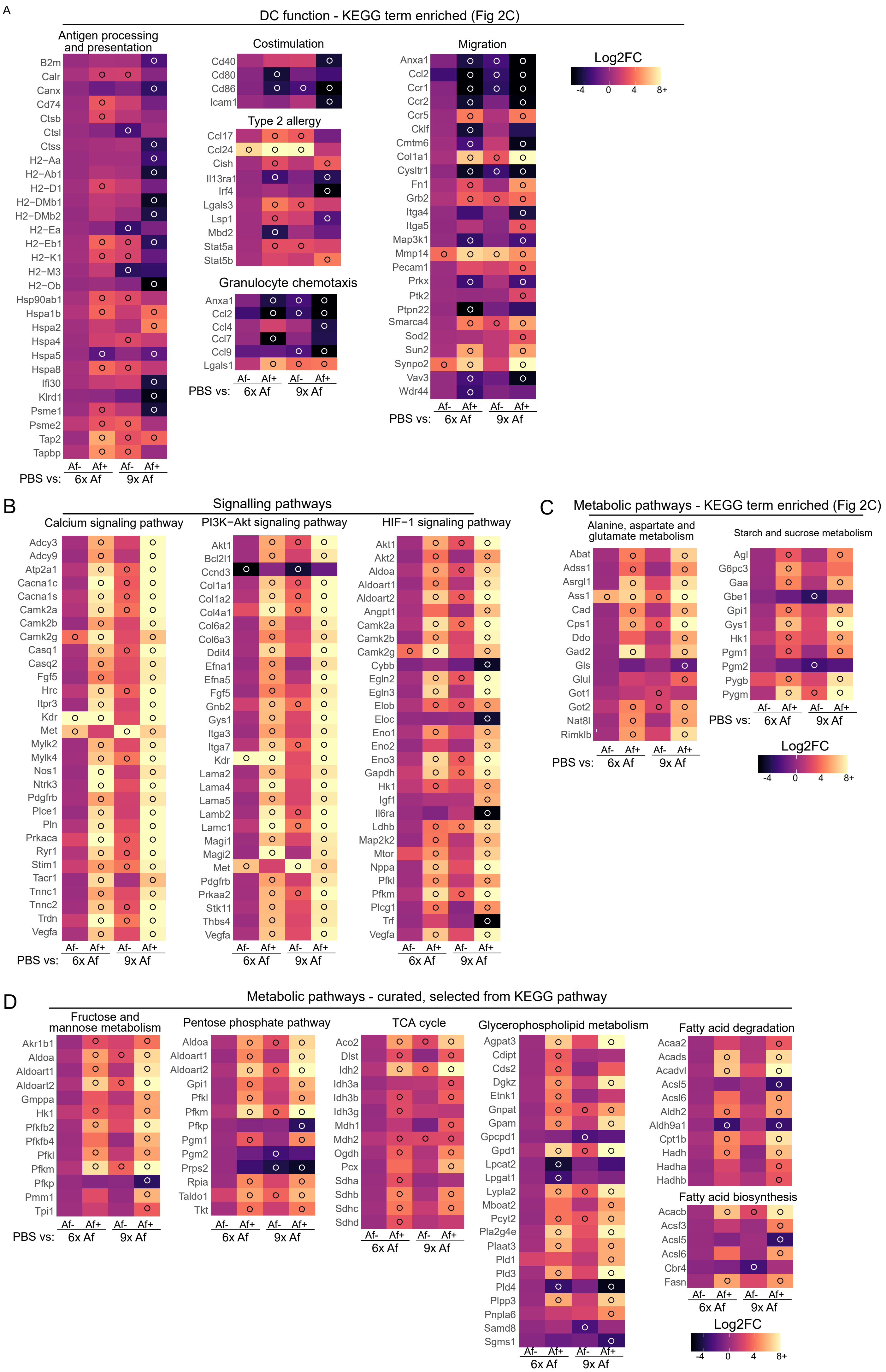


Supp. Figure 8

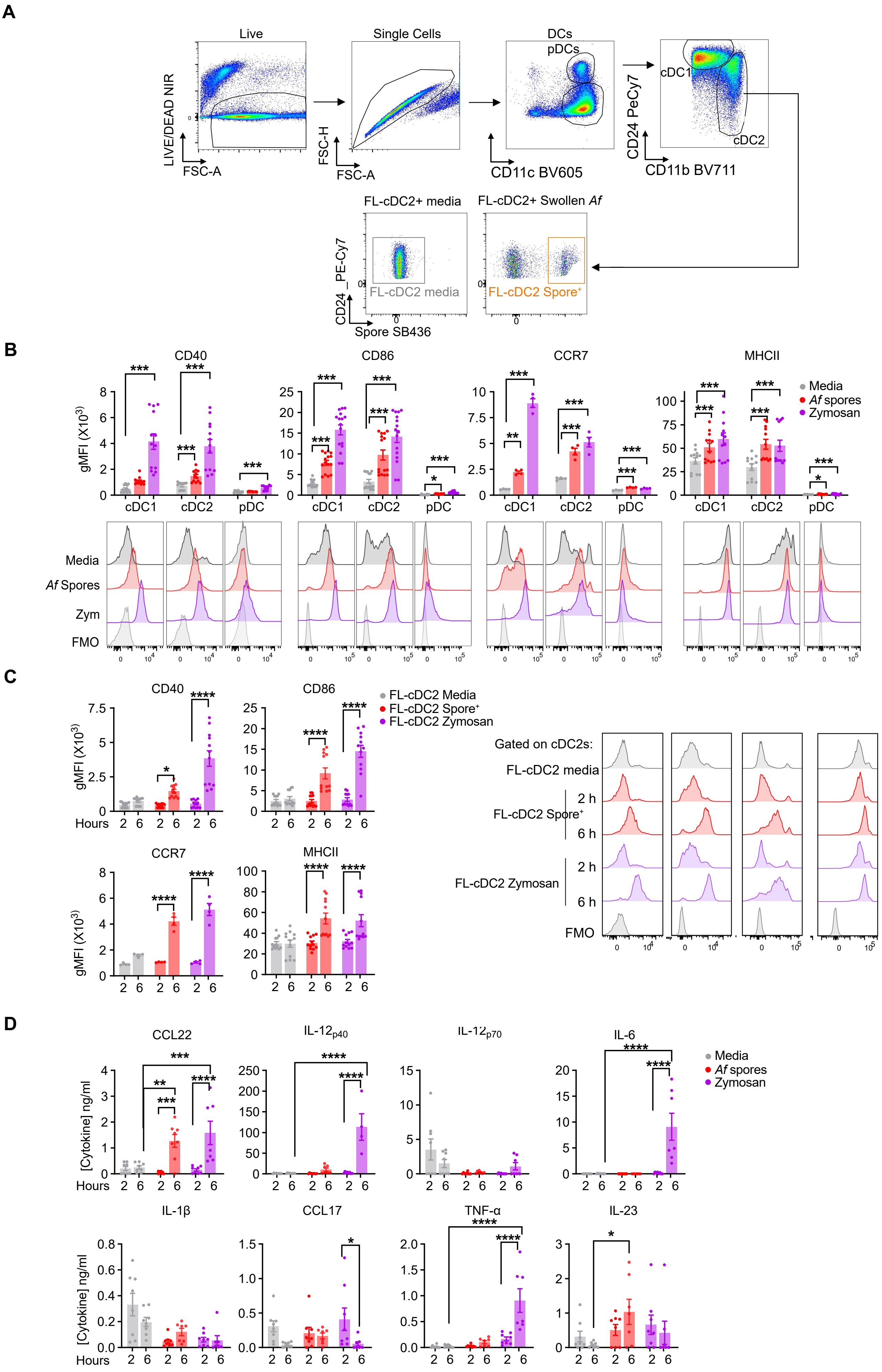


Supp. Figure 9

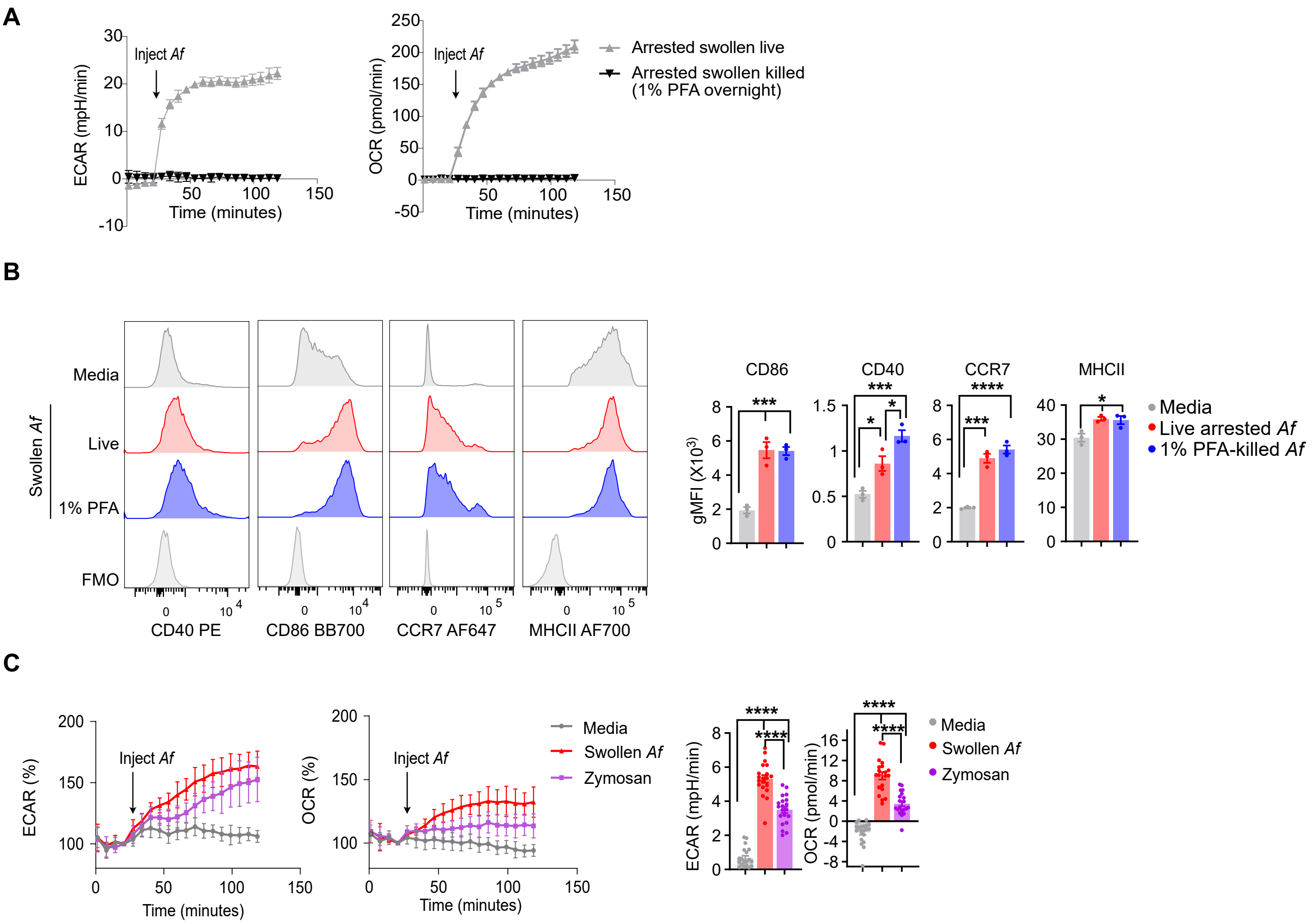


Supp. Figure 10

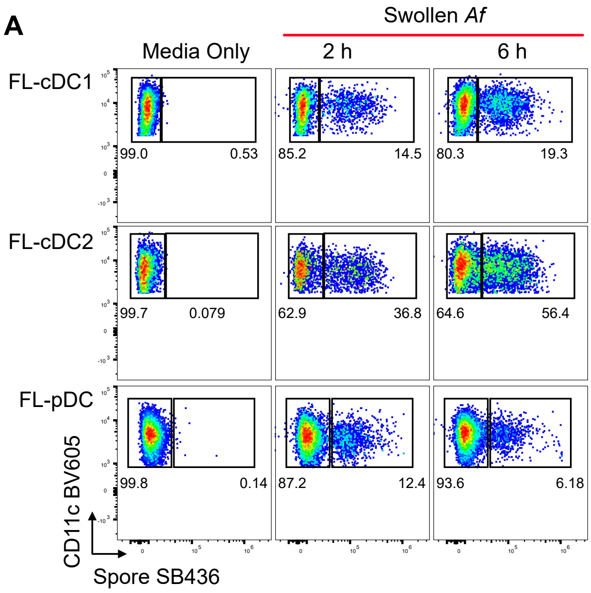


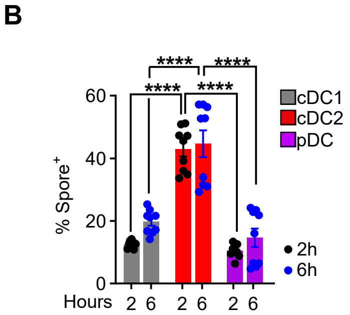


Supp. Figure 11

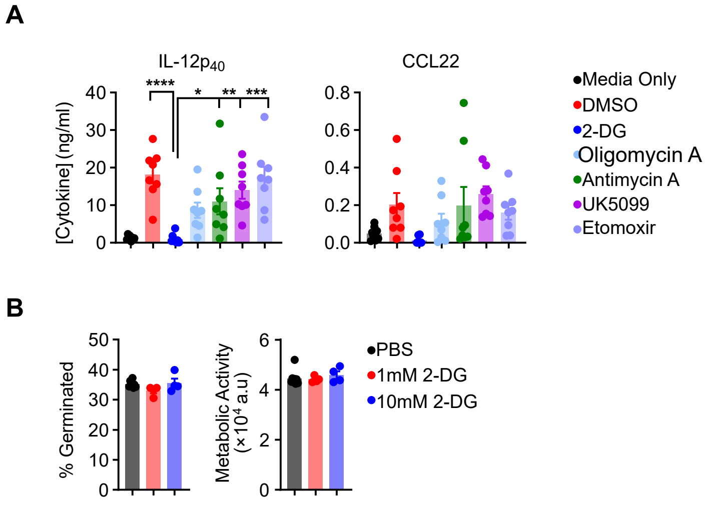


Supp. Figure 12

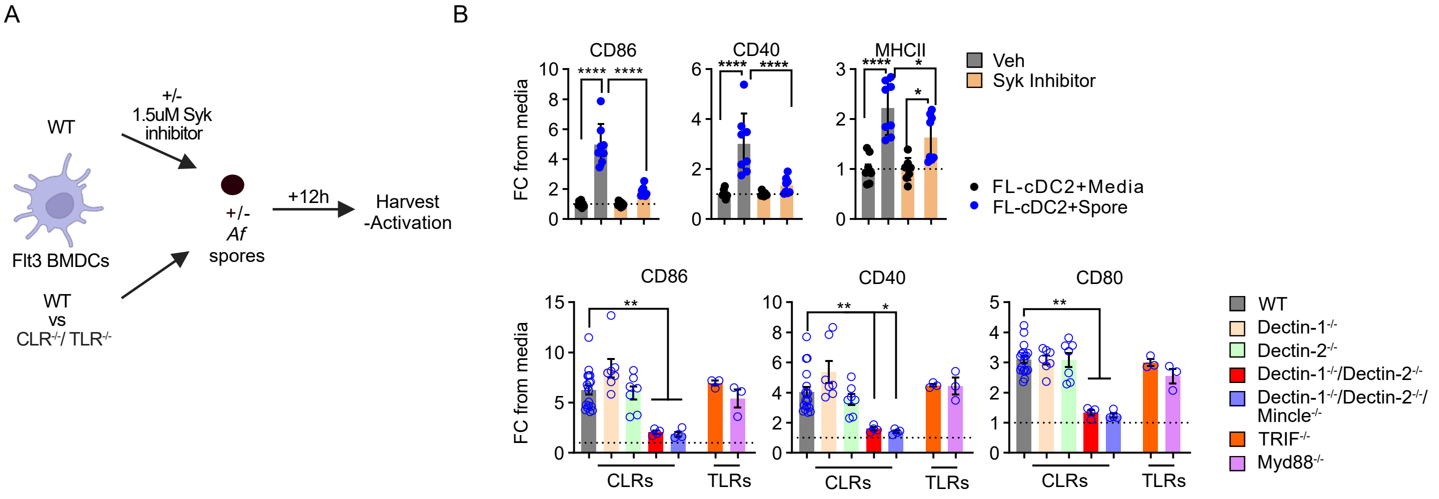


Supp. Figure 13

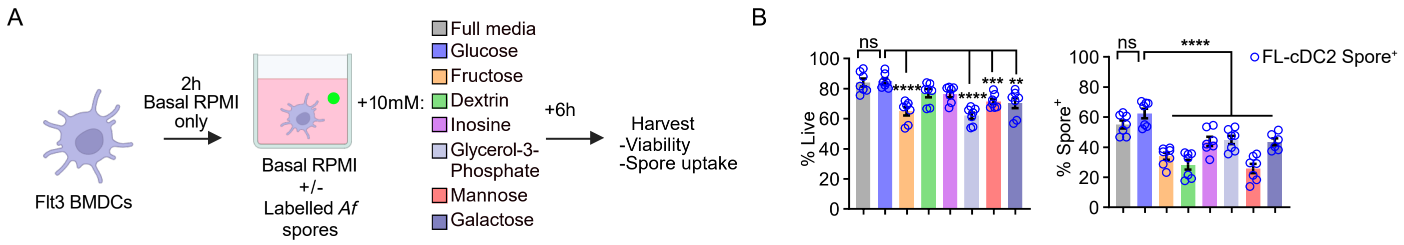


Supp. Figure 14

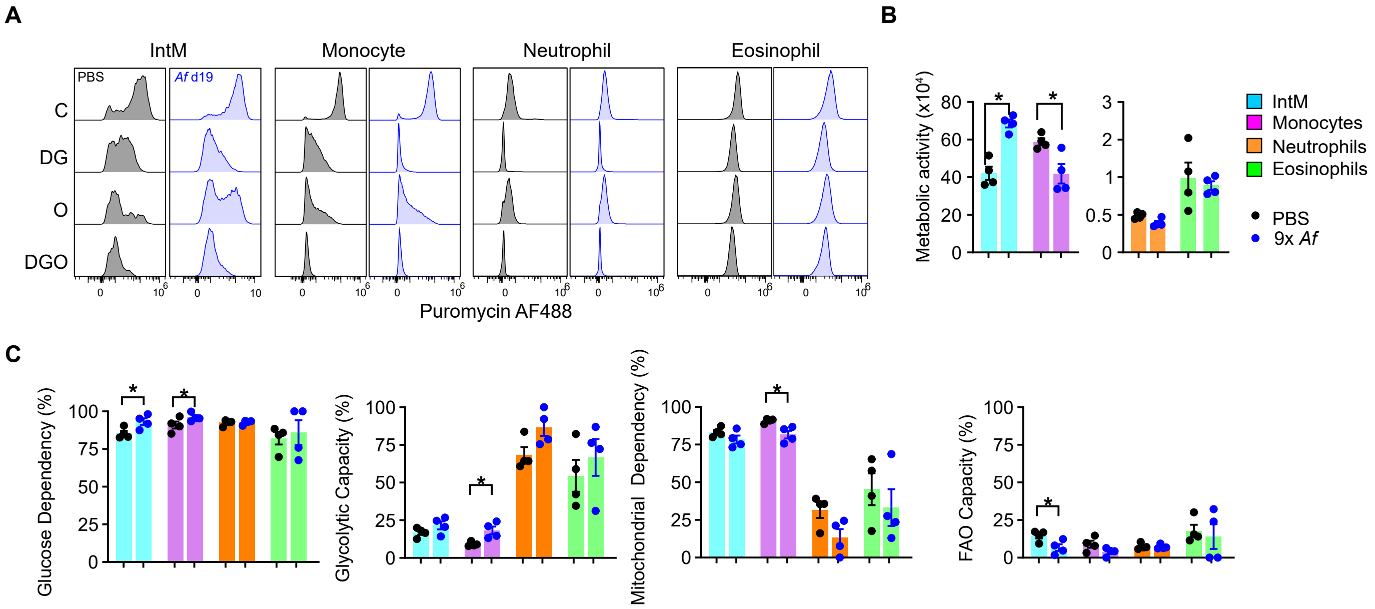


Supp. Figure 15

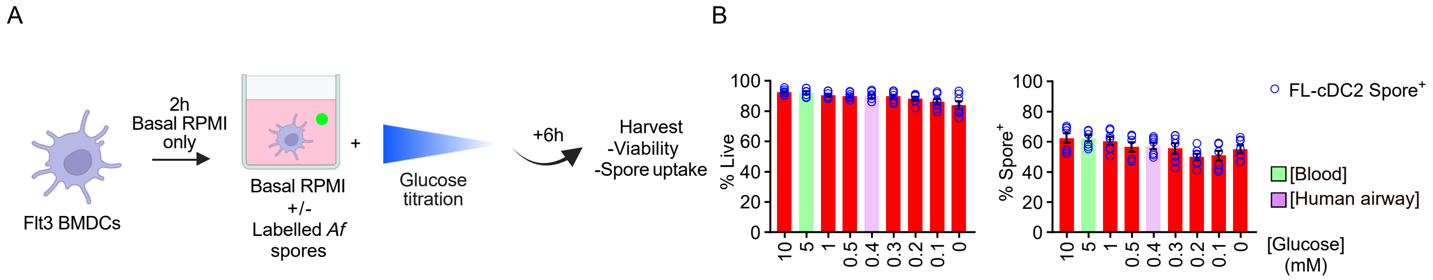


Supp. Figure 16

**Supplementary Table 1: List of antibodies used for flow cytometry and microscopy.**

| Target | Clone | Conjugate | Panel | Titration | Manufacturer |
|---|---|---|---|---|---|
| **SiglecF** | E50-2440 | PE-CF594 | Cell sort | 100 | BD |
| **CX3CR1** | SA011F11 | PE-Dazzle594 | Cell sort | 100 | Biolegend |
| **CD45** | 30-F11 | APC | Confocal Microscopy | 100 | Biolegend |
| **Ep-CAM** | G8.8 | AF488 | Confocal Microscopy | 100 | Biolegend |
| **Pdpn** | Pmab-1 | AF488 | Confocal Microscopy | 100 | Biolegend |
| **CD11c** | N418 | AF647 | FLDC | 200 | Biolegend |
| **MHCII** | M5/114.15.2 | AF700 | FLDC | 1000 | Biolegend |
| **B220** | RA3-6B2 | BV510 | FLDC | 200 | Biolegend |
| **CD11c** | N418 | BV650 | FLDC | 200 | Biolegend |
| **CD11b** | M1/70 | BV711 | FLDC | 1000 | Biolegend |
| **CD45.2** | 104 | FITC | FLDC | 200 | Biolegend |
| **LIVE/DEAD Near IR** | NA | NA | FLDC | 2000 | Thermofisher |
| **CD40** | 3/23 | PE | FLDC | 200 | Biolegend |
| **CD45.1** | A20 | PE | FLDC | 100 | Biolegend |
| **CD24** | M1/69 | PE/Cy7 | FLDC | 800 | Biolegend |
| **CD45** | 30-F11 | BUV395 | Lung | 100 | BD |
| **CD11c** | N418 | BUV496 | Lung | 100 | BD |
| **LIVE/DEAD Blue** | NA | NA | Lung | 500 | Thermofisher |
| **IL-13** | eBio13A | AF488 | Lung cytokine | 200 | Thermofisher |
| **CD4** | RM4-5 | AF700 | Lung cytokine | 100 | Biolegend |
| **CD3** | 17A2 | APC Fire750 | Lung cytokine | 100 | Invitrogen |
| **IL-5** | TRFK5 | BV421 | Lung cytokine | 100 | Biolegend |
| **CD19** | 1D3 | BV480 | Lung cytokine | 100 | BD |
| **CD8** | 53-6.7 | BV510 | Lung cytokine | 100 | Biolegend |
| **TCRgd** | GL3 | BV605 | Lung cytokine | 100 | Biolegend |
| **NK1.1** | PK136 | BV650 | Lung cytokine | 100 | Biolegend |
| **IFNg** | XMG1.2 | BV711 | Lung cytokine | 100 | Biolegend |
| **TCRb** | H57-597 | BV785 | Lung cytokine | 100 | Biolegend |
| **IL-10** | JES5-16E3 | PE | Lung cytokine | 200 | Biolegend |
| **Foxp3** | FJK-16s | PE/Cy5 | Lung cytokine | 100 | Invitrogen |
| **IL-17** | TC11-18H10.1 | PE/Cy7 | Lung cytokine | 100 | Biolegend |
| **IL-4** | 11B11 | PE/Dazzle594 | Lung cytokine | 200 | Biolegend |
| **MHCII** | M5/114.15.2 | AF532 | Lung myeloid | 100 | Invitrogen |
| **CCL19** | NA | AF647 | Lung myeloid | 500 | Protein Foundry |
| **CX3CR1** | SA011F11 | AF647 | Lung myeloid | 100 | Biolegend |
| **CD63** | NVG-2 | AF700 | Lung myeloid | 200 | Biolegend |
| **CD3** | 17A2 | APCeFluor780 | Lung myeloid | 200 | Invitrogen |
| **CD19** | 1D3 | APCeFluor780 | Lung myeloid | 100 | Invitrogen |
| **CD90.2** | 30-H12 | APCeFluor780 | Lung myeloid | 200 | Invitrogen |

| | | | | | |
|---|---|---|---|---|---|
| **CD49b** | DX5 | APCeFluor780 | Lung myeloid | 200 | Invitrogen |
| **NK1.1** | PK136 | APCeFluor780 | Lung myeloid | 200 | Invitrogen |
| **CD26** | 17A2 | BUV737 | Lung myeloid | 100 | BD |
| **SIRPa** | P84 | BUV805 | Lung myeloid | 100 | BD |
| **SiglecF** | E50-2440 | BV480 | Lung myeloid | 400 | BD |
| **XCR1** | ZET | BV510 | Lung myeloid | 100 | Biolegend |
| **Ly6C** | HK1.4 | BV605 | Lung myeloid | 100 | Biolegend |
| **PDCA1** | 927 | BV650 | Lung myeloid | 400 | BD |
| **CD64** | X54-5/7.1 | BV711 | Lung myeloid | 100 | Biolegend |
| **Ly6G** | 1A8 | BV750 | Lung myeloid | 400 | Biolegend |
| **PDL2** | TY25 | BV786 | Lung myeloid | 200 | BD |
| **CX3CR1** | SA011F11 | PacBlue | Lung myeloid | 100 | Biolegend |
| **CD209a** | MMD3 | PE | Lung myeloid | 200 | Biolegend |
| **CD40** | 3/23 | PE/Cy5 | Lung myeloid | 100 | Biolegend |
| **CD301b** | URA-1 | PE/Cy7 | Lung myeloid | 400 | Biolegend |
| **MertK** | 2B10C42 | PE/Dazzle594 | Lung myeloid | 100 | Biolegend |
| **CD11b** | M1/70 | PerCP | Lung myeloid | 100 | Biolegend |
| **Spores** | NA | GFP | Lung non-immune | NA | NA |
| **Ter119** | TER119 | APCeFluor780 | Lung panels | 200 | Invitrogen |
| **Streptavidin** | NA | FITC | Spores | 100 | Biolegend |
| **Streptavidin** | NA | SB436 | Spores | 1200 | Invitrogen |
| **Streptavidin** | NA | BV421 | Spores | 100 | Biolegend |
| **Streptavidin** | NA | AF488 | Spores | 100 | Biolegend |
| **Puromycin** | R4743L-E8 | AF488 | Various | 500 | Gift of R. Arguello |
| **CD11b** | M1/70 | BB700 | Various | 800 | BD |

**Supplementary Table 2. List of antibodies utilised in ELISA.**

| Target | Conjugation | Clone | Manufacturer |
|---|---|---|---|
| **IL-1B** | Purified | - | R&D Duoset |
| **IL-6** | Purified | MP5-20F3 | Biolegend |
| **IL-12p40** | Purified | C15.6 | Biolegend |
| **IL-12p70** | Purified | 9A5 | R&D Duoset |
| **IL-23** | Purified | MMP19B2 | Biolegend |
| **TNFa** | Purified | - | R&D Duoset |
| **CCL17** | Purified | - | R&D Duoset |
| **CCL22** | Purified | - | R&D Duoset |
| **IL-1B** | Biotinylated | - | R&D Duoset |
| **IL-6** | Biotinylated | MP5-32C11 | Biolegend |
| **IL-12p40/70/IL-23** | Biotinylated | C17.8 | Biolegend |
| **TNFa** | Biotinylated | - | R&D Duoset |
| **CCL17** | Biotinylated | - | R&D Duoset |
| **CCL22** | Biotinylated | - | R&D Duoset |
